# Brain endothelial antigen presentation detains CD8^+^ T cells at the blood-brain barrier leading to its breakdown

**DOI:** 10.1101/2021.10.12.464035

**Authors:** Sidar Aydin, Javier Pareja, Vivianne M. Schallenberg, Armelle Klopstein, Thomas Gruber, Nicolas Page, Elisa Kaba, Urban Deutsch, Aaron J. Johnson, Mirjam Schenk, Doron Merkler, Britta Engelhardt

## Abstract

Blood-brain barrier (BBB) breakdown and immune cell infiltration into the central nervous system (CNS) are early hallmarks of multiple sclerosis (MS). High numbers of CD8^+^ T cells are found in MS lesions and antigen (Ag)-presentation at the BBB was proposed to promote CD8^+^ T-cell entry into the CNS. Employing live cell imaging and primary mouse brain microvascular endothelial cells (pMBMECs) as *in vitro* model of the BBB and a mouse model of CNS autoimmunity, we here show that pMBMECs process and present antigens leading to effector CD8^+^ T-cell differentiation. Under physiological flow, endothelial Ag-presentation prohibited CD8^+^ T-cell crawling and diapedesis leading to pMBMEC apoptosis. Reduced motility of Ag-specific CD8^+^ T cells was also observed in CNS microvessels in neuroinflammation *in vivo.* Luminal MHC class I Ag-presentation at the BBB thus prohibits CD8^+^ T-cell entry into the CNS and rather triggers CD8^+^ T cell mediated focal BBB breakdown.

## Introduction

Multiple sclerosis (MS) is a neuroinflammatory disease of the central nervous system (CNS) characterized by blood-brain barrier (BBB) breakdown, immune cell infiltration and demyelination^1^. Accumulating evidence points to a key role for CD8^+^ T cells in MS pathogenesis. The MHC class I allele HLA-A*0301 is associated with increased risk of developing MS^2^. In active MS lesions CD8^+^ T cells outnumber CD4^+^ T cells and oligoclonal expansion of CD8^+^ T cells in MS lesions and cerebrospinal fluid (CSF) suggests they recognize CNS antigens (summarized in^3^). CD8^+^ T cells are furthermore associated with axonal damage as their cytolytic granules polarize towards demyelinated axons^4^. Finally, clinical trials underscore the contribution of lymphocyte subsets other than CD4^+^ T cells to MS pathogenesis^5, 6, 7^.

Immune cell recruitment into the CNS is controlled by the endothelial blood-brain barrier (BBB) (summarized in^8^). Our present knowledge of the molecular mechanisms involved in T-cell trafficking into the CNS is largely based on CD4^+^ T cells, while the mechanisms used by CD8^+^ T cells to enter the CNS in MS is rudimentary. There is emerging evidence that CD8^+^ T cells use molecular mechanisms distinct from those of CD4^+^ T cells to cross the BBB. Following their arrest, CD8^+^ T cells probe and readily cross the BBB endothelium preferentially via a transcellular route, while CD4^+^ T cells rather crawl on the BBB endothelium prior to their diapedesis^8^. Also, although both, CD4 and CD8^+^ T cells use α4β1-integrins to arrest on endothelial vascular cell adhesion molecule (VCAM)-1, only CD8^+^ but not CD4^+^ T cells engage the junctional adhesion molecule (JAM)-B to cross the BBB^8^. While the migration of CD4^+^ T cells across the BBB does not require Ag-specific mechanisms^9^, the migration of CNS autoantigen-specific CD8^+^ T cells across the BBB was suggested to rely on the luminal expression of major histocompatibility complex (MHC) class I^10^. This would imply that antigens released from cellular sources of the CNS parenchyma cross the glia limitans to subsequently be taken up by brain endothelial cells from the abluminal side, processed and cross-presented to circulating CD8^+^ T cells on the luminal side.

This prompted us to study if BBB endothelial cells can (cross)-present luminally and abluminally available exogenous antigens on their luminal surface on MHC class I molecules and if this would facilitate their migration across the BBB *in vitro* and *in vivo.* Our observations show that brain endothelial cells can take up, process and present exogenous antigens on MHC-class I molecules. However, MHC-class I restricted Ag-presentation by the BBB rather induces a stop signal for CD8^+^ T cells recognizing their cognate Ag on the luminal side of the BBB, eventually leading to endothelial cell apoptosis and focal BBB breakdown, which is a critical hallmark of MS pathogenesis.

## Results

### Primary mouse brain microvascular endothelial cells (pMBMECs) present Ag to naïve CD8^+^ T cells

Ag-presentation by professional antigen presenting cells (APCs) to CD8^+^ T cells requires the engagement of the peptide-loaded MHC class I complex with the TCR of the CD8^+^ T cell. Co-stimulation via CD80 and CD86 further initiates the activation of naïve CD8^+^ T cells, while PD-L1 serves as check-point inhibitor regulating T-cell activation. To explore if primary brain microvascular endothelial cells (pMBMECs) are equipped to present Ag we first investigated if pMBMECs express the basic molecular components required for Ag presentation. Immunofluorescence (IF) stainings on non-stimulated pMBMECs showed weak cell surface staining for MHC class I but not for CD80, CD86 or PD-L1. At 24 hours of stimulation with TNF-α/IFN-γ, pMBMECs stained positive for MHC class I and PD-L1 but still not for CD80 or CD86 (Figure 1A). 48 hours of TNF-α/IFN-γ stimulation further increased IF staining for MHC class I and PD-L1 on pMBMECs with CD80 and CD86 still not detectable (Figure 1A). We therefore assessed expression of CD80 and CD86 also at the mRNA level by quantitative real-time PCR (qRT-PCR) in non-stimulated and 5 hours TNF-α/IFN-γ stimulated pMBMECs. While high C_T_ values confirmed low expression of CD80 and CD86 in pMBMECs, TNF-α/IFN-γ stimulated pMBMECs did show a 4.33-fold and 3.52-fold upregulated mRNA expression for CD80 and CD86, respectively, when compared to non-stimulated pMBMECs (Figure 1B, 1C). Cytokine stimulated pMBMECs thus express MHC class I but rather low levels of the costimulatory molecules CD80 and CD86, and high levels of the co-inhibitory molecule PD-L1.

**Figure 1.**
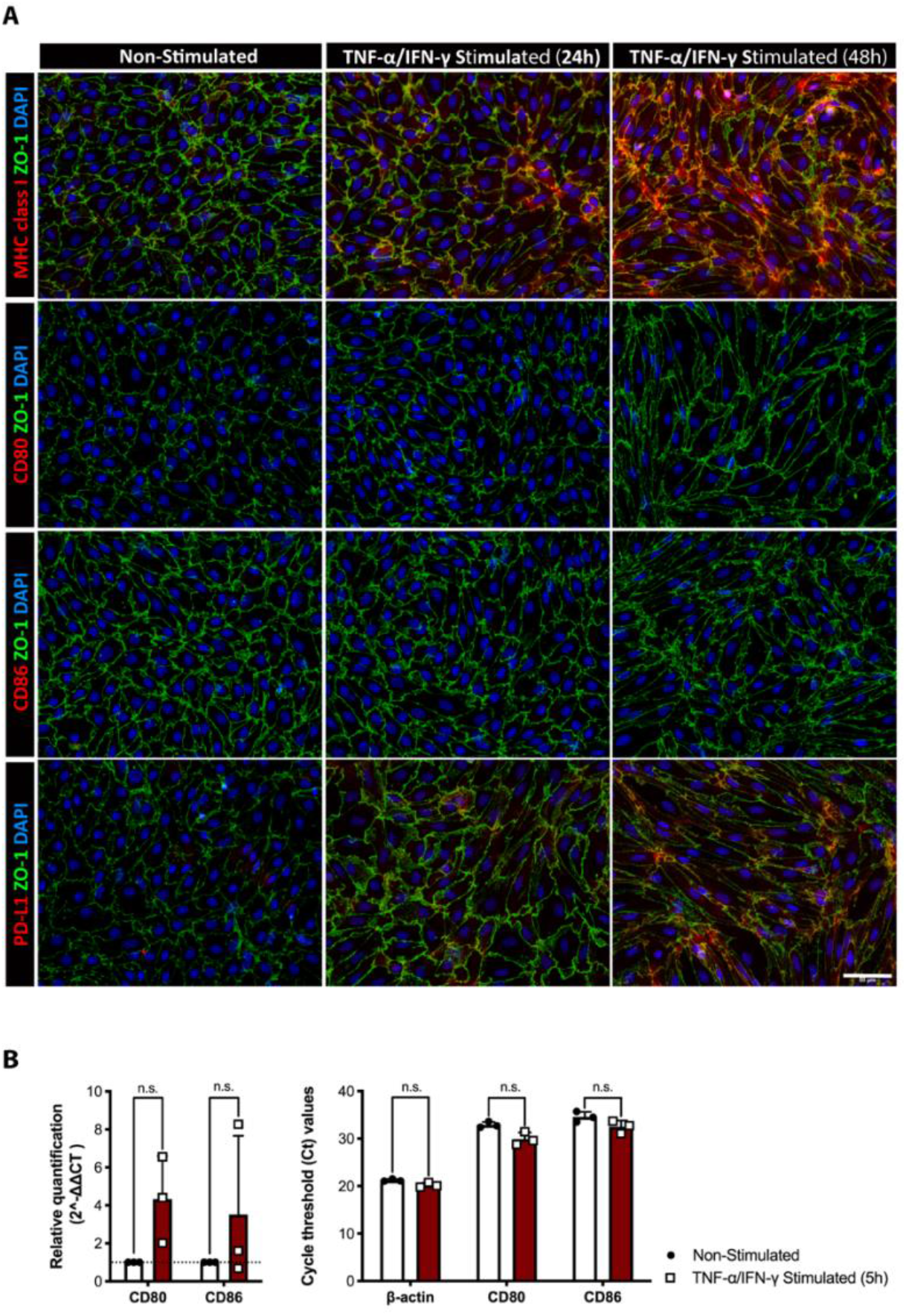
Primary mouse brain microvascular endothelial cells (pMBMECs) can present Ag to the naïve CD8^+^ T cells upon inflammation. **(A)** Immunofluorescence staining of pMBMECs from WT C57BL/6J mice for MHC class I, CD80, CD86 and PD-L1 (red), endothelial cell junctions (ZO-1, green), nuclei (DAPI, blue). pMBMECs were either not stimulated or stimulated with TNF-α and IFN-γ-for 24 and 48 hours. Scale bar = 50 μm. The staining is representative for 5 individual experiments. **(B)** Relative gene expression of CD80 and CD86 in samples from unstimulated and TNF-α/IFN-γ stimulated (5 hours) pMBMECs from WT C57BL/6J mice was assessed by qRT-PCR. For each gene, technical duplicates or triplicates from 3 individual experiments were measured. Relative quantification is represented by the 2^-ΔΔCt^ value (the graph on the left-hand side). The change in the average of C_T_ values of target genes upon stimulation is demonstrated in the bar plots (the graph on the right-hand side). Data were analyzed using non-parametric Mann-Whitney U test *p<0.05, **p<0.01, ***p<0.001, ****p<0.0001.

To explore if this molecular makeup allows TNF-α/IFN-γ stimulated pMBMECs to prime naïve CD8^+^ T cells we co-cultured naïve OT-I T cells for 24 hours with TNF-α/IFN-γ stimulated and antigen (Ag)-peptide pulsed pMBMECs. Peptide-pulsed bone marrow derived dendritic cells (BMDCs) served as professional Ag-presenting cell (APC) control. OT-I cells co-cultured with SIINFEKL-pulsed pMBMECs but not with VSV-control peptide or no peptide-pulsed pMBMECs were readily activated as shown by their enhanced cell surface expression of the early activation protein CD69, the interleukin-2 receptor α-chain CD25 and the adhesion molecule CD44 accompanied with the loss of cell surface expression of cell adhesion molecule CD62L (Figure 2A). Also, B2M^-/-^ pMBMECs, lacking functional MHC class I expression, did not induce any OT-I T cell activation irrespective of their peptide pulsation (Figure 2A). As expected, SIINFEKL-but not VSV-peptide pulsed BMDCs induced activation of the naïve OT-I cells (Figure 2A). Collectively, these results underline that pMBMECs can prime CD8^+^ T cells in an MHC class I dependent manner.

**Figure 2.**
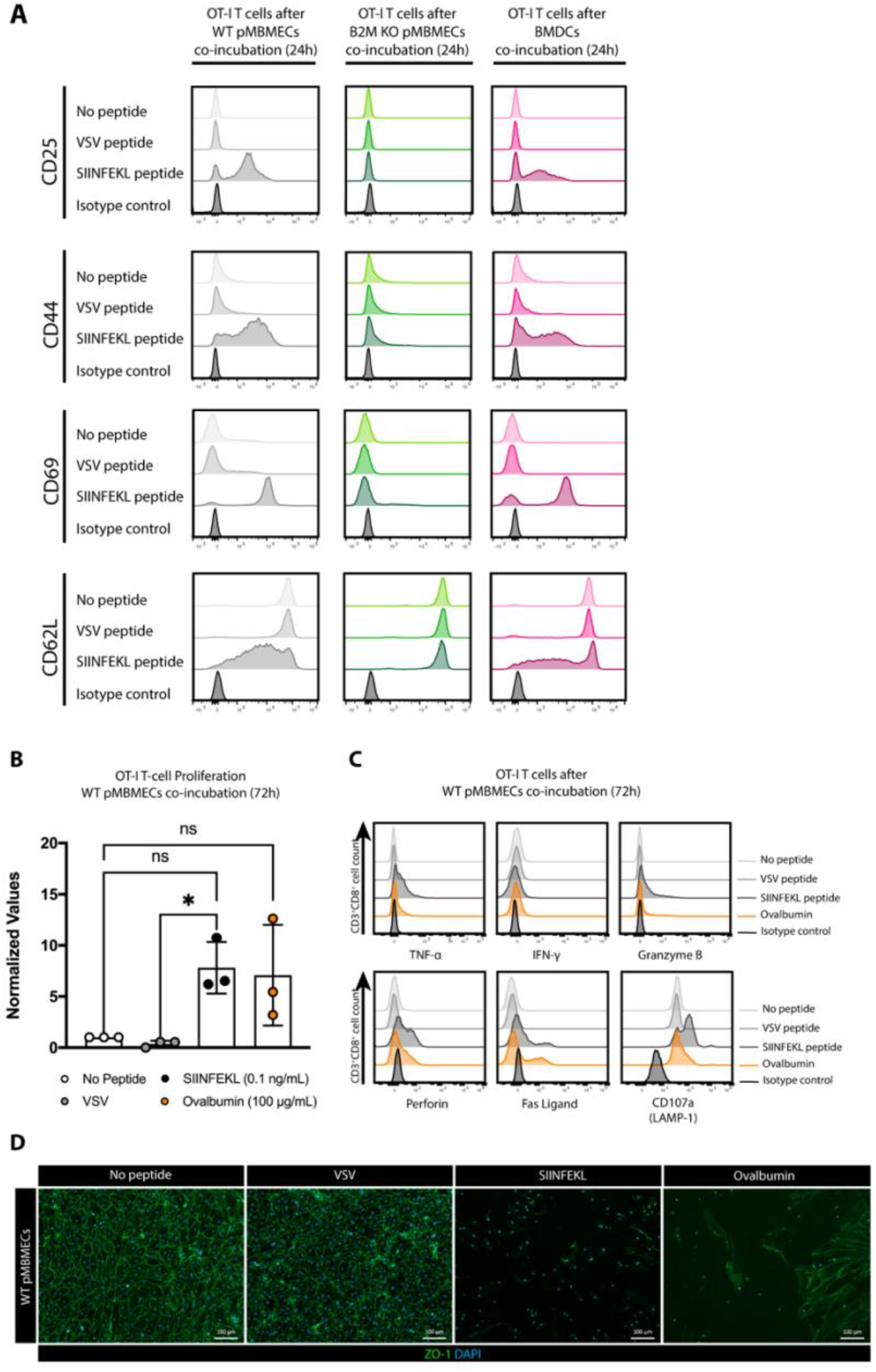
Luminal Ag-presentation by pMBMECs elicit the activation and the proliferation of naïve OT-I cells and initiate the disruption of the barrier *in vitro*. **(A)** TNF-α/IFN-γ stimulated pMBMECs from WT or B2M mice, lacking the functional expression of MHC class I, were pulsed for 30 minutes with the ovalbumin peptide SIINFEKL (0.1 ng/mL). pMBMECs were co-cultured for 24 hours with purified naïve OT-I T cells isolated from OT-I Rag-1^-/-^ C57BL/6J mice. Peptide-pulsed BMDCs from WT C57BL/6J mice were used as professional APC positive control. Multicolor flow cytometry analysis of naïve OT-I cells after 24 hours of co-culture with TNF-α/IFN-γ stimulated pMBMECs from WT (Grey) or B2M^-/-^ mice (Green) or activated BMDCs (Pink) from WT C57BL/6J mice in the absence of peptide or in the presence of VSV-or SIINFEKL peptide for each condition is shown. Histograms depict cell surface staining of CD3^+^CD8^+^ OT-I cells for CD25, CD44, CD69 and CD62L after co-incubation with pMBMECs. Isotype control for each marker in each condition is shown at the bottom of each plot. The data is representative of 3 individual experiments. **(B)** The assessment of naïve OT-I cell proliferation by BrdU-incorporation assay after 72 hours of co-culture with TNF-α/IFN-γ stimulated WT pMBMECs in the absence of peptide or in the presence of VSV-peptide, SIINFEKL or incubation with full-length ovalbumin (OVA) is shown. The data are pooled from 3 individual experiments with technical duplicates or triplicates. Data were analyzed using one-way ANOVA (*p<0.05, **p<0.01, ***p<0.001, ****p<0.0001) and are shown as mean ± SD normalized to the condition without peptide pulsation. **(C)** Flow cytometry analysis of naïve OT-I cells after 72 hours of coculture with WT pMBMECs in the absence of peptide or in the presence of VSV-peptide, SIINFEKL or fulllength OVA. Histograms show cell surface staining of CD3^+^CD8^+^ OT-I cells from the co-culture for TNF-α, IFN-γ, granzyme B, perforin, Fas ligand and CD107a (LAMP-1). Isotype control for each marker in each condition is shown at the bottom of each plot. The data is representative of 3 individual experiments. **(D)** Immunofluorescence staining of TNF-α/IFN-γ stimulated WT pMBMECs for endothelial cell junctions (ZO-1, green), nuclei (DAPI, blue) after 72 hours of co-culture with naïve OT-I cells. Loss of junctional integrity and endothelial cells is visible after co-culturing of naïve OT-I cells with SIINFEKL-pulsed or OVA-loaded WT pMBMECs. The data is representative of 3 individual experiments. Scale bar = 100 μm.

Therefore, we next asked if TNF-α/IFN-γ stimulated WT pMBMECs pulsed with the SIINFEKL peptide can also induce OT-I proliferation following their activation. 72 hours of co-incubation of naïve OT-I T cells with SIINFEKL-pulsed but not with no peptide or VSV -peptide pulsed pMBMECs induced T cell proliferation as assessed by incorporation of bromodeoxyuridine (BrdU) into the OT-I cells (Figure 2B). Pulsation of pMBMECs with the peptides fitting the MHC class I peptide binding site does not allow to explore if pMBMECs can take up and degrade exogenous protein antigens, load peptides on their MHC class I molecules and transport them to the surface for Ag-presentation. To explore this, we co-incubated of TNF-α/IFN-γ stimulated WT pMBMECs with the full-length ovalbumin protein and naïve OT-I cells and observed also under these conditions OT-I cell proliferation to a level comparable to SIINFEKL-pulsed pMBMECs. Activated pMBMECs can therefore process exogenous protein antigens for presentation on MHC class I molecules inducing CD8^+^ T cell activation and proliferation. To finally explore if brain endothelial cell Ag-presentation allowed for induction of CD8^+^ T cell effector functions we determined induction of effector molecule expression in OT-I cells after co-incubation with TNF-α/IFN-γ stimulated pMBMECs pulsed with either SIINFEKL or the VSV-control peptide or loaded with OVA. Flow cytometry analysis for the cytokines TNF-α and IFN-γ, cytotoxic granule molecules perforin and granzyme B (GrB), lysosome-associated membrane glycoprotein (LAMP)-1 and the apoptosis-inducing transmembrane protein fas ligand (FasL) showed already at 48 hours of co-incubation upregulation of IFN-γ and GrB in OT-I cells co-incubated with SIINFEKL-pulsed or OVA-loaded pMBMECs (data not shown). After 72 hours of co-incubation upregulation of TNF-α, perforin, GrB, LAMP-1 and FasL could be detected in OT-I cells co-incubated with SIINFEKL-pulsed pMBMECs and in a similar manner with OVA-loaded pMBMECs (Figure 2C). In contrast, no peptide-pulsed and VSV-peptide pulsed pMBMECs did not induce any upregulation of cytotoxic effector molecules in OT-I cells (Figure 2C). Expression of cytotoxic effector molecules lead to the gain of effector functions in OT-I cells leading to the disruption of the pMBMEC monolayers as determined by IF staining for endothelial junctions at 72 hours of co-incubation (Figure 2D). To understand the dynamics of OT-I cell – pMBMEC interactions in the presence or absence of cognate Ag over time we finally investigated the interaction of tdTomato expressing naïve OT-I cells with unpulsed or SIINFEKL-pulsed pMBMECs isolated from VE-cadherin-GFP reporter mice by in vitro live cell imaging. Observing the interaction of naïve OT-I cells with the pMBMECs over 72 hours showed that the number of tdTomato^+^ OT-I cells co-incubated with non-pulsed pMBMECs decreased over time while the pMBMEC monolayer remained intact (Supplementary Video 1). In contrast, OT-I cells co-incubated with SIINFEKL-pulsed pMBMECs increased in size and initiated disruption of the pMBMEC monolayer as visualized by disruption of the VE-cadherin-GFP labeled junctions (Supplementary Video 1). At the same time SIINFEKL-pulsed pMBMECs without OT-I cells remained intact for the entire observation time.

Taken together our data show that cytokine stimulated pMBMECs can take up, process and present exogenous Ag to naïve OT-I CD8^+^ T cells, leading to their priming, activation and proliferation and induction of effector functions causing destruction of the pMBMEC monolayer.

### Endothelial Ag-presentation impairs the crawling of naïve CD8^+^ T cells under physiological flow *in vitro*

When interacting with the endothelial vascular wall in the physiological context T cells encounter shear forces within the blood stream. To understand if brain endothelial Ag presentation to naïve CD8^+^ T cells can also be observed under shear flow, we studied the interaction of naïve OT-I cells with pMBMECs under physiological flow by *in vitro* live cell imaging. TNF-α/IFN-γ stimulated WT or B2M^-/-^ pMBMECs were pulsed with either SIINFEKL or VSV peptides or left unpulsed. Upon superfusion over pMBMECs, low and comparable numbers of naïve OT-I cells arrested on the inflamed endothelium irrespective of the presence or absence of cognate Ag and functional MHC class I, underscoring that initial arrest of naïve OT-I cells on the pMBMEC monolayer under physiological flow was independent of Ag-recognition (Figure 3A, Supplementary Video 2). However, while after transient arrest a significant fraction of OT-I cells readily detached from the pMBMEC monolayers under most conditions this was not observed on SIINFEKL-pulsed WT pMBMECs suggesting that endothelial Ag-presentation prohibited OT-I cell detachment (Figure 3B, Supplementary Video 2). To further explore how endothelial Ag-presentation affects the post-arrest behavior of naïve OT-I cells on pMBMECs under flow we quantified the dynamic interaction of the individual OT-I cells on the pMBMEC monolayers by performing a visual frame-by-frame offline analysis of the time-lapse videos. This analysis showed that in the absence of endothelial Ag-presentation, the majority of naïve OT-I cells crawled over the pMBMEC monolayer with a minority showing a probing behavior (Figure 3B). Diapedesis of naïve OT-I cells across the pMBMEC monolayer was however rarely observed. In contrast, on SIINFEKL -pulsed pMBMECs, most naïve OT-I cells remained stationary and actively probed with cellular protrusions the pMBMEC surface while only few OT-I cells crawled over the pMBMEC monolayer (Figure 3B). The few OT-I cells observed crawling over SIINFEKL-pulsed pMBMEC showed a reduced crawling speed and crawling distance when compared to OT-I cells crawling over pMBMECs lacking presentation of the cognate Ag (Figure 3C, D).

**Figure 3.**
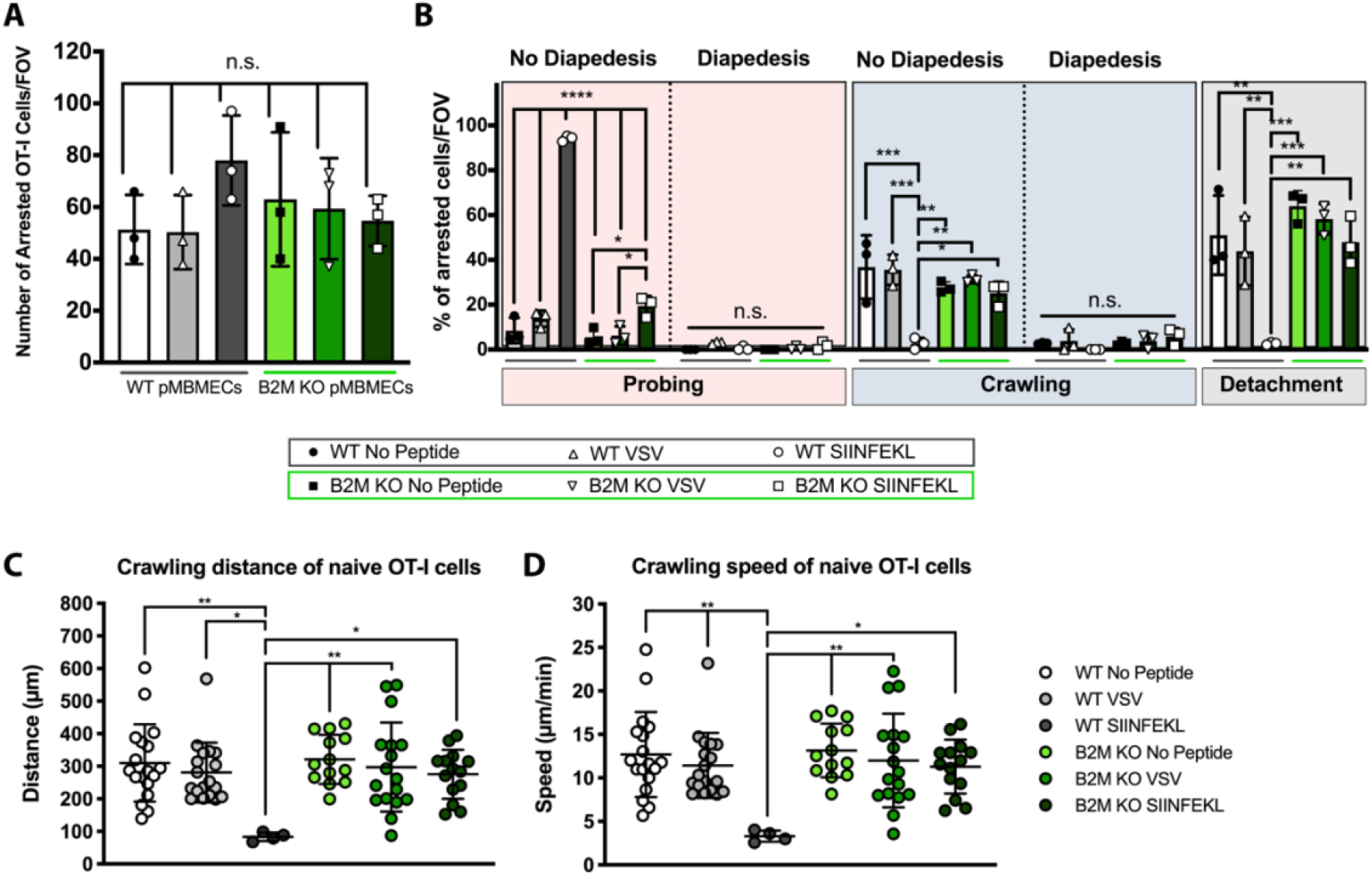
Naïve OT-I T cells display impaired crawling on inflamed pMBMECs upon recognition of cognate Ag in an MHC class I dependent fashion under physiological flow *in vitro*. **(A)** Number of arrested naïve OT-I cells per FOV (872 x 654 μm) on TNF-α/IFN-γ stimulated WT pMBMECs without peptide pulsing (white bar), with VSV peptide pulsing (grey bar), SIINFEKL peptide pulsing (black bar) and on B2M^-/-^ pMBMECs without peptide pulsing (light green bar), with VSV peptide pulsing (green bar), SIINFEKL peptide pulsing (dark green bar). A total of three individual experiments were conducted. **(B)** Quantification of post-arrest dynamic behavior of naïve OT-I cells on WT and B2M^-/-^ pMBMECs during the 30 min recording time. The number of arrested OT-I T cells for each condition was set to 100% and the behavioral categories are shown as fraction thereof. Presence or absence of complete diapedesis is indicated for each migratory behavioral category. The percentage of the partial diapedesis after both probing and crawling for each condition were below 2% without any significant difference and are not shown in the graph. Data is shown as mean ± SD from three individual experiments. **(C,D)** Crawling distance in μm **(C)** and speed **(D)** in μm/min of naïve OT-I cells on TNF-α/IFN-γ stimulated WT and B2M^-/-^ pMBMECs are shown. Each dot represents one crawling track of one naïve OT-I cell. Values are pooled from three individual expeirments and shown as mean ±SD. For each condition 4-18 cells were tracked. Data were analyzed using repeated measure ANOVA with Tukey post-test *p<0.05, **p<0.01, ***p<0.001, ****p<0.0001.

To our surprise, the higher avidity of naïve OT-I cell interaction with SIINFEKL-pulsed pMBMECs combined with their probing behavior did not result in their diapedesis across the pMBMEC monolayer (Figure 3B). Taken together, our observations thus show that naïve CD8^+^ T cells can arrest on brain endothelial cells in an Ag-independent manner under physiological flow. Endothelial Ag-presentation does however influence their post-arrest behavior seemingly by inducing a stop signal that prohibits naïve CD8^+^ T cell detachment from and crawling on the pMBMEC monolayer and rather induces their continuous probing of the luminal pMBMEC.

### BBB endothelial cells can process Ags from the abluminal side to prime naïve CD8^+^ T cells in an MHC class I dependent manner at the luminal side

The barrier properties of the BBB do not allow for free diffusions of molecules. pMBMECs have previously been shown to establish a tight barrier towards the diffusion of small molecules^11^. To explore if BBB endothelial cells could present CNS derived Ag on their luminal side we next investigated if pMBMECs can take up and process exogenous Ag from their abluminal side and present this Ag on MHC class I on their luminal surface to naïve CD8^+^ T cells. WT pMBMECs were grown on filter inserts to confluency allowing to mimic luminal and abluminal compartments of the *in vitro* BBB model. Increasing concentrations of SIINFEKL or OVA were introduced from the abluminal compartment after luminal stimulation of pMBMECs with TNF-α/IFN-γ and naïve OT-I cells were co-incubated with pMBMECs for 24 and 72 hours on the luminal surface. Abluminal exposure of WT pMBMECs with SIINFEKL peptide and OVA protein induced naïve OT-I cell activation as shown by their up-regulation of CD69, CD25 and CD44, and loss of cell surface expression of CD62L (Figure 4A). In contrast, OT-I T cells co-cultured with pMBMECs in the absence of added abluminal Ag remained naïve (Figure 4A). Following their activation, OT-I cells proliferated (Figure 4B) and differentiated into effector cells leading to the disruption of the pMBMEC monolayers (Figure 4C).

**Figure 4.**
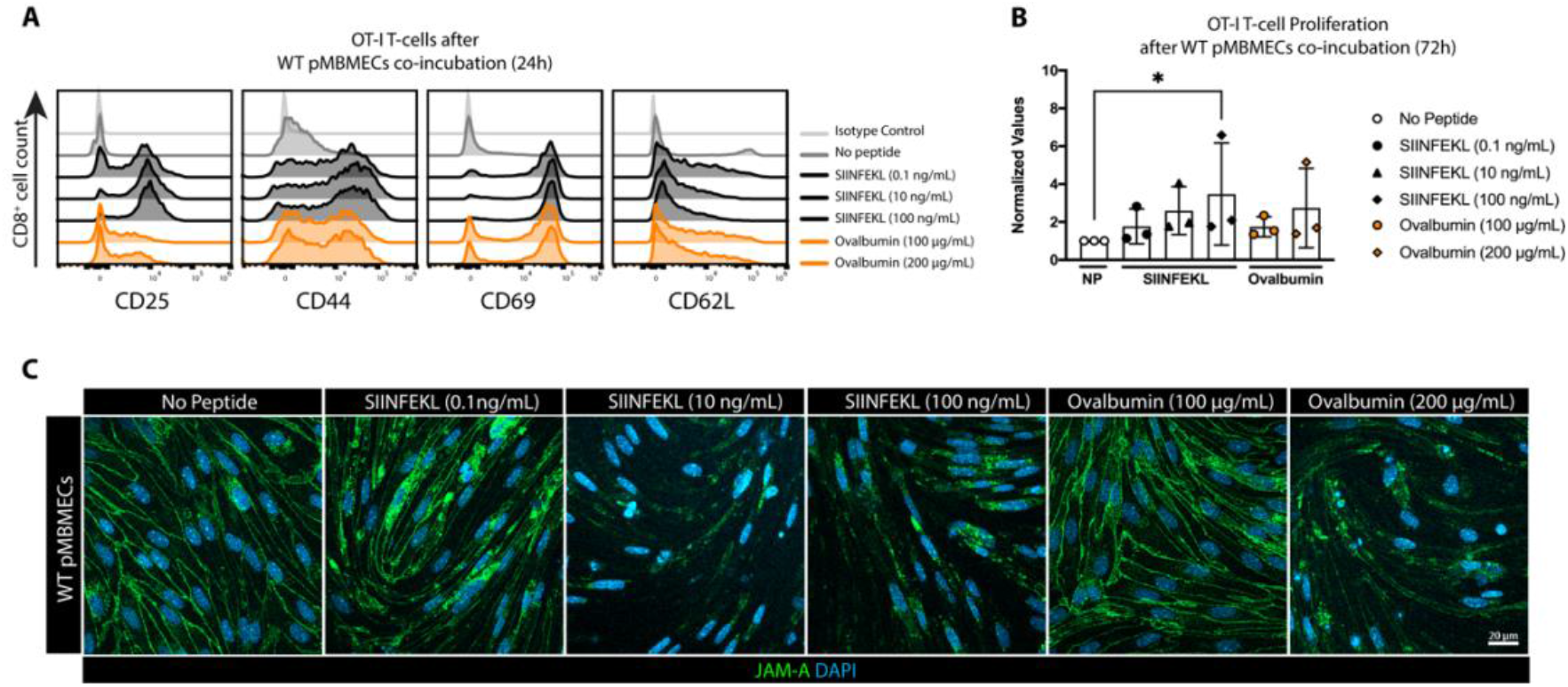
pMBMECs cross-present abluminal Ags to naïve OT-I cells on their luminal side and induce their differentiation into effector cells *in vitro*. **(A)** Flow cytometry analysis of naïve OT-I cells for activation markers after 24 hours of co-culture with TNF-α/IFN-γ stimulated pMBMECs from WT C57BL/6J mice on the Transwell filters with abluminal presence of VSV-peptide, SIINFEKL or OVA (SIINFEKL: 0.1 ng/mL, OVA: 100 μg/mL). Histogram plots show cell surface staining of CD3^+^CD8^+^ naïve OT-I cells after co-incubation for CD25, CD44, CD69 and CD62L. Isotype control for each marker in each condition is shown at the bottom of each plot. The data is representative of 3 individual experiments. **(B)** The assessment of naïve OT-I cell proliferation by BrdU-proliferation assay after 72 hours of co-culture with TNF-α/IFN-γ stimulated pMBMECs from WT C57BL/6J mice on the Transwell filters in the absence of Ag or presence of abluminal SIINFEKL or OVA with increasing concentrations (SIINFEKL: 0.1 ng/mL, 10 ng/mL, 100 ng/mL; OVA: 100 μg/mL, 200 μg/mL). The data are pooled from 3 individual experiments with technical duplicates or triplicates. Data were analyzed using one-way ANOVA (*p<0.05, **p<0.01, ***p<0.001, ****p<0.0001) and are shown as mean ± SD normalized to the condition without peptide pulsation. **(C)** Immunofluorescence staining of TNF-α/IFN-γ stimulated pMBMECs from WT C57BL/6J mice for endothelial cell junctions (ZO-1, green), nuclei (DAPI, blue) after 72 hours of co-culture with naïve OT-I cells on the Transwell filters with the absence of an Ag or abluminal presence of SIINFEKL or OVA with increasing concentrations (SIINFEKL: 0.1 ng/mL, 10 ng/mL, 100 ng/mL; OVA: 100 μg/mL, 200 μg/mL). Scale bar = 20 μm. The data is representative of 3 individual experiments.

In summary, pMBMECs can process exogenous Ag from their abluminal side for presentation on MHC class I molecules on their luminal surface allowing for naïve CD8^+^ T cell priming and activation and differentiation into effector CD8 T cells leading to focal BBB breakdown.

### Brain endothelial Ag-presentation stops effector CD8^+^ T cell migration and initiates CD8^+^ T cell-mediated BBB breakdown

Our observations showed that differentiation of effector CD8^+^ T cells upon MHC class I restricted Ag-recognition on pMBMECs ultimately resulted in disruption of the brain endothelial monolayer. Therefore, we next asked how Ag-presentation by pMBMECs would affect the interaction with effector CD8^+^ T cells under physiological shear flow. We therefore studied the interaction of *in vitro* activated effector OT-I cells with pMBMECs under physiological flow by *in vitro* live cell imaging. TNF-α/IFN-γ stimulated WT or B2M^-/-^ pMBMECs were pulsed with either SIINFEKL or VSV peptides or left unpulsed. Activated OT-I cells arrested on pMBMECs with approximately 7-fold higher numbers than naïve OT-I cells and their initial arrest was not affected by the presence or absence of cognate Ag or functional MHC class I (Figure 5A, Supplementary Video 3). In the absence of SIINFEKL, effector OT-I cells readily crossed the WT or B2M^-/-^ pMBMEC monolayer following their probing or crawling (Figure 5B, Suppl. Video 3). In contrast, on SIINFEKL pulsed WT pMBMECs OT-I cells showed significantly reduced crawling followed by diapedesis and increased probing behavior without diapedesis (Figure 5B, Supplementary Video 3). Also, among the small portion of activated OT-I cells observed to continuously crawl on the pMBMECs, both, the crawling speed and distance were reduced on SIINFEKL-pulsed WT pMBMECs when compared to all other conditions (Figure 5C, D). Thus, recognition of their cognate Ag on the pMBMECs initiated a stop signal for OT-I cells as observed by their increased probing behavior and their reduced diapedesis upon crawling across the pMBMEC monolayer. The Ag-specific arrest of activated OT-I cells on the pMBMEC monolayer under flow resulted in rapid disruption of the pMBMEC monolayer observed already at 20 minutes after initiation of the interaction (Figure 5E, Supplementary Video 4).

**Figure 5.**
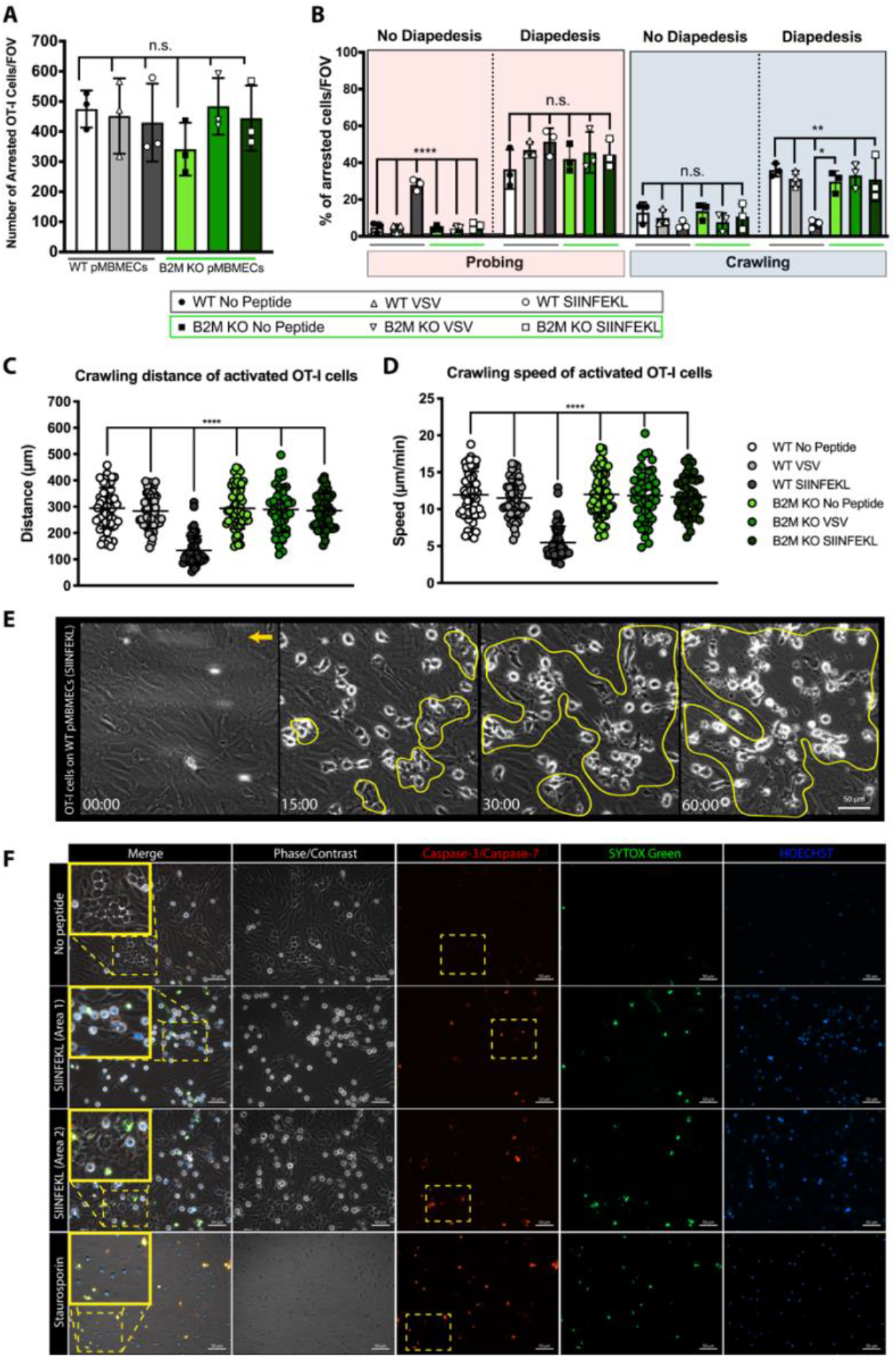
Ag -presentation by the pMBMECs stops the migration of effector OT-I cells and initiates the barrier breakdown in a granzyme B dependent fashion under physiological shear flow *in vitro*. **(A)** Number of arrested *in vitro* activated effector OT-I cells per FOV (872 x 654 μm) on TNF-α/IFN-γ stimulated WT pMBMECs without peptide pulsing (white bar), with VSV peptide pulsing (grey bar), SIINFEKL peptide pulsing (black bar) and on B2M^-/-^ pMBMECs without peptide pulsing (light green bar), with VSV peptide pulsing (green bar), SIINFEKL peptide pulsing (dark green bar). A total of three individual experiments were conducted. Data is shown as mean ± SD. **(B)** Post-arrest dynamic behavior of effector OT-I cells on WT and B2M^-/-^ pMBMECs during 30 min of recording time. The behavioral categories are demonstrated as percentage of categorized effector OT-I cells for each condition on pMBMECs. Presence or absence of complete and partial diapedesis is indicated for each migratory behavioral category. The percentage of T cell detachment was below 2% for each condition without any significant difference and is not shown in the graph. Data is shown as mean ± SD from three individual experiments. **(C,D)** Crawling distance in μm **(C)** and speed **(D)** in μm/min of effector OT-I cells on TNF-α/IFN-γ stimulated WT and B2M^-/-^ pMBMECs are shown. Each dot represents one crawling track of one effector OT-I cell. For each condition 44-60 cells were tracked. Values were pooled from three individual experiments and shown as mean ±SD. Data were analyzed using repeated measure ANOVA with Tukey post-test *p<0.05, **p<0.01, ***p<0.001, ****p<0.0001. **(E)** Representative image sequence of effector OT-I cell induced killing of SIINFEKL pulsed pMBEMCs under physiological flow during 60 minutes of recording time. The area of endothelial cell killing is marked with yellow shapes. The orange arrow indicates the direction of the flow. Scale bar = 20 μm. The data is representative of 3 individual experiments. **(F)** Live staining of pMBMECs with Image-iT™ LIVE Red Poly Caspases Detection Kit following 60 minutes of intercations with activated OT-I cells physiological flow. Phase/contrast imaging allowed us to determine the disrupted areas of pMBMECs. SYTOX^®^ Green nucleic acid stain was used to differentiate the live and dead cells, whereas HOECHST staining was used to detect the cell nuclei. Grey: phase constrast imaging to visualize the pMBMEC monolayer; Red: Caspase-3/-7 staining to detect apopotosis, Green: Sytox Green to detect cell membrane damage, Blue: HOECHST staining to viusalize cell nuclei. Scale bar = 50 μm. The data is representative of 3 individual experiments.

To understand if the OT-I cell mediated destruction of the pMBMEC monolayer is induced by the cytotoxic activity of OT-I cells leading to apoptosis of the pMBMECs we next explored the role of the cytotoxic effector proteins GrB, which was observed to be upregulated in OT-I cells upon *in vitro* priming (Supplementary Figure 2). We crossed OT-I mice with GrB deficient C57BL/6J mice (OT-I GrB^-/-^) allowing for subsequent side-by-side comparison of WT and GrB^-/-^ effector OT-I cells with TNF-α/IFN-γ stimulated, SIINFEKL pulsed or unpulsed VE-cadherin-GFP pMBMECs under physiological flow by live cell imaging.

The newly generated OT-I GrB^-/-^ cells did now show any impairment of their migratory behavior, as we did not observe any difference in the number of arrested cells and their post-arrest behavior between OT-I and OT-I GrB^-/-^ cells (data not shown). However, while upon cognate Ag recognition, effector OT-I cells readily disrupted the pMBMEC monolayer this was not observed during the interactions of effector OT-I GrB^-/-^ cells with pMBMEC under physiological flow (Supplementary Video 5). Thus, activated OT-I cells can recognize their Ag on MHC class I under physiological flow, leading to the release of cytotoxic granules in brain endothelial cell death probably by apoptosis.

To verify if pMBMECs die by OT-I induced apoptosis, unpulsed or SIINFEKL-pulsed pMBMECs were live stained for active caspase activity after one hour of interaction with effector OT-I cells using the fluorescent inhibitor of caspases (FLICA™) methodology. Positive Caspase-3/Caspase-7 staining in combination with membrane damage was observed in SIINFEKL-pulsed pMBMECs but not in unpulsed pMBMECs upon co-culture with effector OT-I cells (Figure 5F). Importantly, different regions of SIINFEKL-pulsed pMBMECs showed a heterogeneous pattern in monolayer disruption. While some areas of the pMBMEC monolayers had detached following cell apoptosis, remaining pMBMECs showed signs of apoptosis and membrane damage prior to their detachment (Figure 5F).

Priming of naïve CD8^+^ T cells into effector or memory CD8^+^ T cells with a specific Ag is determined by the alterations of numerous cell surface and intracellular molecules^12^. It is, thus, conceivable that *in vitro* priming of CD8^+^ T cells may induce effector functions that do not fully resemble those observed in CD8^+^ T cells upon *in vivo* activation^13^. To verify if our *in vitro* primed CD8^+^ T cells are a suitable model, we next isolated CD8^+^ T cells from tdTomato^+^ OT-I C57BL/6J mice and activated the OT-I cells in *in vitro* and *in vivo. In vitro* priming of naïve tdTomato^+^ OT-I cells was performed by a co-culture with professional APCs and the cognate peptide SIINFEKL, whereas for *in vivo* priming naïve tdTomato^+^ OT-I cells were injected into WT C57BL/6J recipient mice 24 hours prior to their infection with full length OVA expressing-lymphocytic choriomeningitis virus (LCMV). *In vivo* and *in vitro* primed tdTomato^+^ OT-I cells were stained for CD25, CD69, CD44, CD62L, TNF-α, IFN-γ, GrB, perforin and LAMP-1 for side-by-side flow cytometry analysis. *In vitro* and *in vivo* activated OT-I cells were found to express comparable cell surface levels for CD62L, CD44 and CD69, whereas *in vivo* activated OT-I cells showed lower levels of CD25 (Figure 7A, Supplementary Figure 4). At the same time, *in vivo* activated OT-I cells showed higher levels of IFN-γ, comparable levels of TNF-α and lower levels of GrB and perforin in comparison to *in vitro* activated OT-I cells. While *in vitro* activated OT-I cells showed a homogenous and intense LAMP-1 immunostaining, *in vivo* activated OT-I cells divided into two subsets with high and low staining intensities (Figure 7A). As there are subtle differences in the activation profile of *in vitro* versus *in vivo* activated OT-I cells, we next aimed to understand if this would impact on the dynamic behavior of OT-I cells on pMBMECs under physiological shear flow. We, therefore, studied the behavior of *in vivo* activated OT-I cells with TNF-α/IFN-γ stimulated and SIINFEKL-pulsed or unpulsed pMBMECs under physiological flow by *in vitro* live cell imaging. While *in vivo* activated OT-I cells arrested in lower numbers on the pMBMEC monolayers when compared to *in vitro* activated OT-I cells, they did not show any difference in their post-arrest behavior on pMBMECs under flow. On SIINFEKL-pulsed pMBMECs *in vivo* primed OT-I cells showed reduced crawling behavior and diapedesis when compared to unpulsed pMBMECs as already observed with *in vitro* primed OT-I cells (Supplementary Figure 4A). Moreover, *in vivo* primed OT-I cells also mediated apoptosis of SIINFEKL-pulsed pMBMECs (Supplementary Figure 4B). Overall *in vivo* activated OT-I effector cells allowed to recapitulate our observations made with the *in vitro* activated OT-I cells underscoring that upon recognition of their cognate Ag on brain endothelial cells effector CD8^+^ T cells stop migrating and induce BBB disruption.

Albeit rare, pMBMEC cultures always contain few pericytes even after their negative selection^14^. The Ag presentation capacity of pericytes has not been studied extensively^15^. Therefore, to not overlook a potential Ag-presentation event by the few pericytes in the pMBMECs culture, we asked if pericytes rather than pMBMECs would initiate the MHC-class I restricted Ag-presentation to CD8^+^ T cells in pMBMEC monolayers. To this end, we isolated pMBMECs from NG2-DsRed C57BL/6J mice to visualize the remaining pericytes in pMBMEC cultures. TNF-α/IFN-γ stimulated, SIINFEKL-pulsed or unpulsed-pMBMEC - effector OT-I T-cell interactions were observed under physiological flow by live cell imaging as described above. We found that effector OT-I cells continued crawling over the pericytes in the SIINFEKL pulsed-cultures and did neither specifically stop on pericytes nor preferentially induce pericyte death (Supplementary Fig 3).

Taken together, these results demonstrate that MHC class I restricted Ag recognition by effector CD8^+^ T cells on the BBB endothelium is not involved in the initial CD8^+^ T cell arrest but post arrest serves as a stop signal for CD8^+^ T cells on the BBB. This leads to reduced CD8^+^ T cell diapedesis and instead to CD8^+^ T cell mediated-, GrB dependent endothelial apoptosis, causing the disruption of the endothelial cell monolayer and barrier breakdown *in vitro* under flow.

### Endothelial Ag-cross-presentation facilitates arrest and probing of naïve and effector CD8^+^ T cells on the inflamed BBB *in vivo*

Finally, we asked how brain endothelial cell antigen presentation of CNS antigens may impact on CD8^+^ T cell interaction with the BBB. To this end, we employed the ODC-OVA mouse, which expresses OVA as a neo-self Ag in oligodendrocytes that is solely visible to CD8^+^ T cells but not to CD4^+^ T cells or B cells^16^. CNS autoimmune inflammation was induced by adoptive transfer of naïve tdTomato^+^ OT-I cells followed 24 hours later by a peripheral infection with LCMV-OVA in ODC-OVA and in WT C57BL/6J control mice. As previously reported 5-6 days following LCMV-OVA infection, ODC-OVA mice exhibited clinical symptoms, accompanied with weight loss, whereas WT controls did not show any clinical disease symptoms^17^.

To verify MHC class I expression on brain endothelial cells *in vivo* we performed immunostainings on brain and spinal cord sections from ODC-OVA and WT C57BL/6J mice for MHC-I and podocalyxin as a vascular marker on day 7 after LCM-OVA infection (Figure 6A and Supplementary Figure 5). MHC class I immunostaining was shown to strongly colocalize with podocalyxin^+^ microvessels in spinal cord and brain grey and white matter in ODC-OVA and WT C57BL/6J mice indicating a homogeneous expression of MHC class I in the CNS microvessels (Figure 6A and Supplementary Figure 5). Positive immunostaining for MHC class I on CNS parenchymal cells was additionally observed in ODC-OVA but not C57BL/6 control mice, underscoring the ongoing neuroinflammation in the ODC-OVA mice (Figure 6A and Supplementary Figure 5).

**Figure 6.**
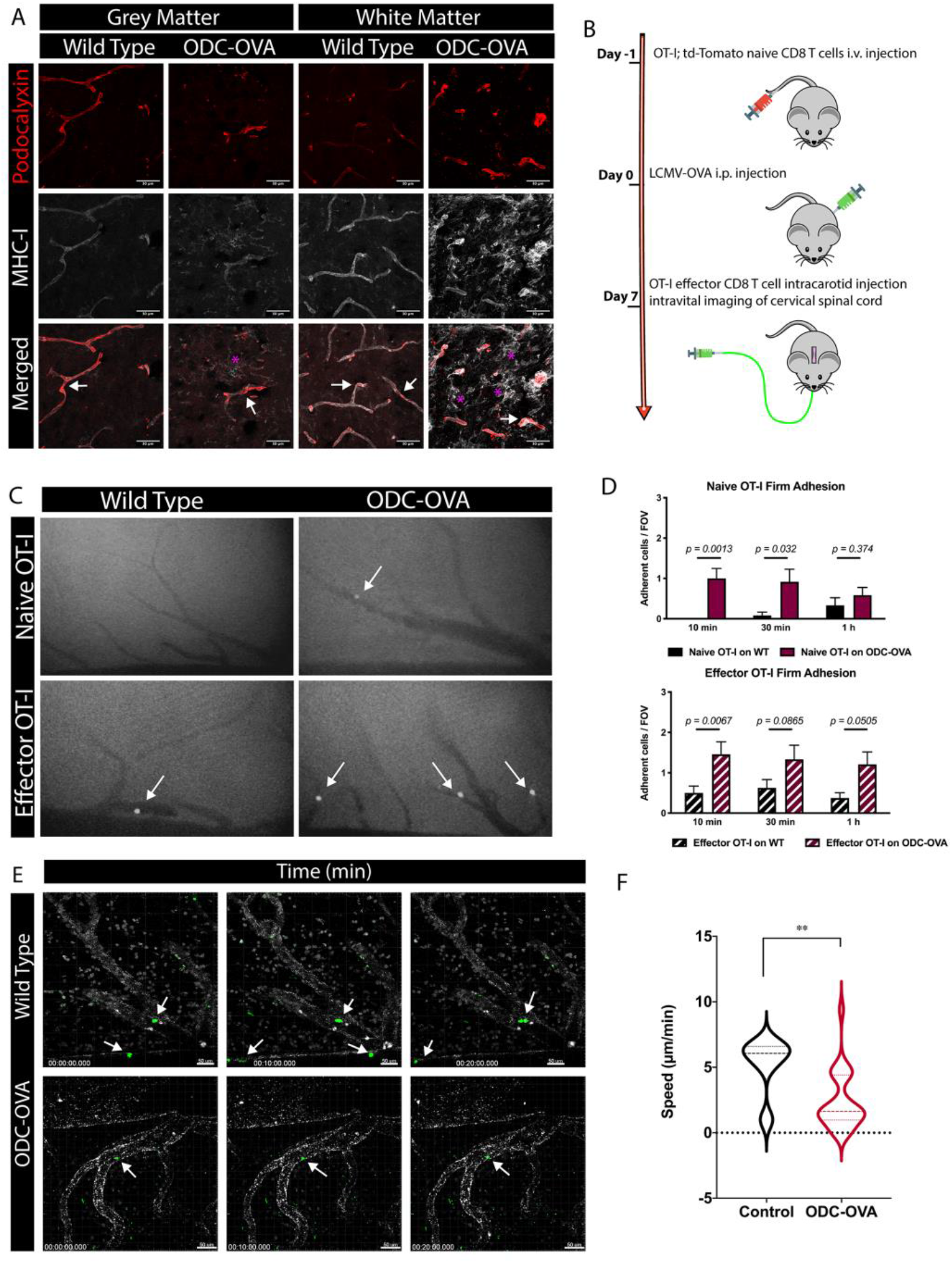
CD8^+^ T cells show an increased arrest coefficient as well as lower general motility on the inflamed BBB *in vivo*. **(A)** Immunofluorescence staining of WT C57BL/6J and ODC-OVA mouse brains on day 7 after LCMV-OVA infection for MHC class I (grey), and the vasculature (podocalyxin, red). Grey matter stainings are displayed in the left panel and white matter stainings in the right panel. White arrows show CNS vessels with signal from both MHC class I and podocalyxin. Purple asterisks point to other CNS resident cells expressing MHC class I molecules on their surface. Scale bar = 30 μm. The data is representative of 5 repetitions from 2 individual experiments. **(B)** Experimental setup of intravital microscopy for the CD8^+^ T cell driven neuroinflammation model. **(C)** Representative images of epifluorescence-IVM imaging of Cell Tracker Green (CMFDA) labeled-naïve and effector OT-I cells adhering in inflamed cervical spinal cord microvessels of WT and ODC-OVA C57BL/6J mice with neuroinflammation 60 minutes post-infusion. White arrows show arrested OT-I cells in cervical spinal cord microvasculature. **(D)** Number of adhered naïve and effector OT-I cells in inflamed cervical spinal cord microvessels of WT and ODC-OVA C57BL6J mice with neuroinflammation at 10, 30 and 60 minutes after infusion of OT-I cells. Data were pooled from imaging of 3 mice/genotype for naïve- and 4 mice/genotype for effector OT-I cells, analyzed using non-parametric Mann-Whitney U test *p<0.05, **p<0.01, ***p<0.001, ****p<0.0001 and are shown as ± SEM. **(E)** Representative images over time of two-photon imaging of CMFDA-labeled effector OT-I CD8^+^ T cells (green) adhered in inflamed cervical spinal cord microvessels (white) of WT or ODC-OVA C57BL/6J mice 60 minutes post-infusion on day 7 after viral infection. The pictures are depicted as maximum intensity projection of 100 μm thick Z-stacks. White arrows point to OT-I effector CD8^+^ T cells. **(F)** Violin plots of the crawling speed (μm/min) of the CMFDA-labelled effector CD8^+^ T cells injected immediately prior to intravital imaging on the luminal spinal cord microvascular wall (a total of 6 cells were analyzed in control mice and 25 in the ODC-OVA mice). Cell tracking was performed with Imaris 9.2 software. Data were analyzed using two-sided unpaired non-parametric T-test. *p<0.05, **p<0.01, ***p<0.001, ****p<0.0001.

To investigate if we could observe Ag-specific differences in the dynamic interaction of CD8^+^ T cells with the BBB *in vivo* according to our observations made *in vitro,* we investigated the interactions of naïve and effector OT-I CD8^+^ T cells with the inflamed BBB endothelium in ODC-OVA mice and LCMV-OVA infected WT C57BL/6J controls. Consecutive to cervical spinal cord window preparation, we systemically injected CMFDA-labelled naïve or effector OT-I cells via a carotid artery catheter and studied their interactions with spinal cord microvessels in real time by epifluorescence intravital microscopy (eIVM) (Figure 6B).

While in WT C57BL/6 mice we hardly observed any interaction of naïve OT-I cells with the spinal cord microvessels, very few naïve OT-I cells showed transient arrest in spinal cord microvessels in the ODC-OVA mice (Figure 6C). When compared to naïve OT-I cells, effector OT-I cells arrested in higher numbers on the inflamed spinal cord microvessels and showed sustained arrests (Figure 6C, D). Observing visibly higher numbers of OT-I cells arresting in spinal cord microvessels of OCD-OVA mice when compared to WT controls (Figure 6C, D) suggests a specific impact of the ongoing neuroinflammation on CD8^+^ T cell interaction with the BBB *in vivo.*

As endothelial Ag-presentation reduced the crawling speed of OT-I cells on pMBMECs *in vitro,* we next asked if such reduced crawling speeds of OT-I cells on the BBB could also be observed specifically in ODC-OVA mice *in vivo.* We performed two-photon intravital microscopy (2P-IVM) of the cervical spinal cord microvessels in ODC-OVA mice and C57BL/6J and VE-Cadherin-GFP C57BL/6 mice as controls on day 7 after LCMV-OVA infection (Figure 6E, Supplementary Videos 6, 7). CMFDA-labelled effector OT-I CD8^+^ T cells were systemically injected via a carotid artery catheter prior to imaging. Comparing the dynamic interaction of these *in vitro* activated effector CD8^+^ T cells with the luminal wall of spinal cord microsvessels in ODC-OVA versus control mice showed a significantly reduced crawling speed of OT-I cells in the ODC-OVA mouse (Figure 6F). Finally, we additionally analyzed the interaction of the still recirculating tdTomato^+^ *in vivo* activated effector OT-I cells with the spinal cord microvessels and observed a similar significant decrease in their crawling speed, their displacement and directionality in ODC-OVA mice when compared to WT control mice (Figure 7B, C).

**Figure 7.**
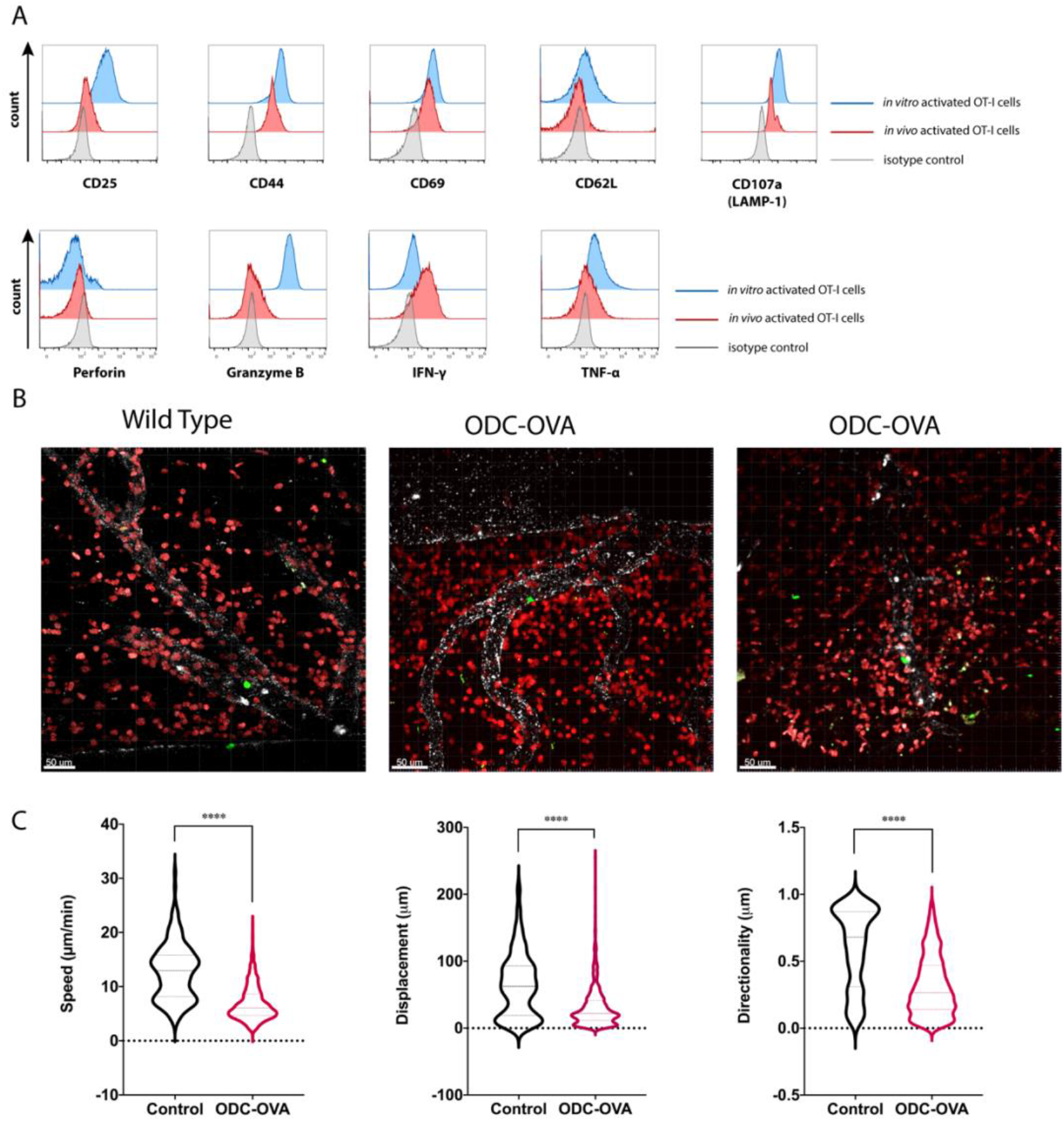
*In vitro* activated CD8^+^ T cells behave comparably to *in vivo* activated CD8^+^ T cells. **(A)** Side-by-side comparison of *in vitro* and *in vivo* activated OT-I cells via flow cytometry. Histograms show flow cytometry staining for CD25, CD44, CD69, CD62L, TNF-α, IFN-γ, GrB, Perforin and CD107a (LAMP-1) of CD3^+^CD8^+^ single-cell gated naïve OT-I cells (Figure 4A) upon their *in vitro* (blue) and *in vivo* (red) activation, while isotype control for each marker is shown with plots in grey. The data is representative of 3 individual experiments. **(B)** Representative images of two-photon imaging of *in vivo* activated tdTomato^+^ effector OT-I cells (red) in inflamed cervical spinal cord microvessels (white) of WT or ODC-OVA C57BL/6J mice on day 7 after LCMV-OVA infection. The pictures are depicted as maximum intensity projection of 100 μm thick Z-stacks. Scale bar = 50 μm. **(C)** General motility measures of *in vivo* activated tDtomato^+^ effector OT-I injected one day prior to LCMV-OVA infection recirculating inside the spinal cord microvessels (196 cells were analyzed in the control mice and 991 in the ODC-OVA mice). Cell tracking was performed with Imaris 9.2 software. Data were analyzed using two-sided un-paired non-parametric T-test. *p<0.05, **p<0.01, ***p<0.001, ****p<0.0001.

Our observations underscore that in ODC-OVA mice with ongoing neuroinflammation, CD8^+^ T cells establish stronger interactions with the BBB endothelial cells resulting in a reduction of CD8^+^ T cell crawling in favor of probing. Taken together, the interaction of CD8^+^ T cells with the BBB in ODC-OVA mice when compared to WT control mice thus resembles CD8^+^ T cell interaction with pMBMECs upon recognition of their cognate Ag in an MHC class I dependent manner.

## Discussion

In the present study we show that BBB endothelial cells can process exogenous protein antigens from both their abluminal and luminal side and present it on their luminal side in an MHC class I dependent manner to CD8^+^ T cells *in vitro.* Effective Ag-dependent activation of naïve T cells is a process usually involving two signals. The first Ag-dependent signal is provided by the engagement of the T-cell receptor with its cognate Ag presented on the MHC molecule on the APC. The second signal is Ag-independent and mediated by co-stimulatory molecules such as CD80 and CD86 on the APC, engaging CD28 on the T cell. The need for a second signal in T cell activation is thought to ensure the maintenance of self-tolerance.

Here we observed that inflamed pMBMECs despite their very low expression of CD80 and CD86, supported priming and activation of naïve CD8^+^ T cells in an MHC class I dependent manner leading to CD8^+^ T cell proliferation and full effector cell differentiation as shown by induction of brain endothelial apoptosis and ultimately rupture of the pMBMEC monolayer. Reports on endothelial expression of CD80 and CD86 have been controversial with some studies showing expression of both co-stimulatory molecules while others unable to detect mRNA or protein expression of these molecules in endothelial cells (summarized in^15^. However, human brain endothelial cells were previously shown to support allogeneic activation and proliferation of naïve CD4^+^ and CD8^+^ T cells in the absence of CD80 and CD86 expression suggesting the existence of mechanisms allowing to bypass this co-stimulatory pathway in T-cell activation. Moreover, some T-cell responses can be induced in mice lacking expression of CD80/CD86 or CD28^18^. In case of endothelial cells, molecules such as CD40L and ICOS-L^19^ or adhesion molecules such as ICAM-1 and VCAM-1 were shown to provide co-stimulatory signals for T-cell activation (summarized in^15^. Thus, on cytokine stimulated pMBMECs low expression of CD80 and CD86 may be compensated by the expression of other co-stimulatory molecules including ICAM-1 and VCAM-1^20^. Alternatively, endothelial MHC class I-peptide complexes may suffice to activate naïve OT I CD8^+^ T cells as previously shown in studies aiming to delineate the minimal requirements for naïve CD8^+^ T cell activation by using APC-free systems^21^. These results suggested that a potent TCR signal can bypass the requirement for co-stimulation. Taking into account that OT-I CD8^+^ T cells recognize the ovalbumin derived peptide SIINFEKL bound to H2K^b^ with high affinity^22^, we therefore chose an experimental setup where we pulsed pMBMECs with very low concentrations of SIINFEKL at 0.5nM and lower to avoid high Ag-density on pMBMECs^21^. Methodology allowing to specifically quantify the peptide-MHC class I complexes on the surface of pMBMECs would allow to measure the Ag density on pMBMECs and the requirement of co-stimulation for naïve CD8 T cell activation by pMBMECs.

An additional observation we made is that inflamed pMBMECs possess the entire machinery allowing for cross-presentation of Ag^23^ as they can take up, process and present exogenously provided ovalbumin on MHC class I molecules on their luminal surface leading to naïve OT-I activation. Our observations are in accordance to previous studies showing that brain endothelial cell crosspresentation of parasite Ags during experimental cerebral malaria induces CD8^+^ T cell mediated BBB dysfunction^24^. In cerebral malaria, brain endothelial cells take up parasite Ags from their luminal side to subsequently cross-present these Ags again on their luminal surface to circulating CD8^+^ T cells. Our present study shows, however, that inflamed brain microvascular endothelial cells can also process exogenous protein Ags provided from their abluminal side and present it on MHC class I molecules to naïve CD8^+^ T cells on their luminal side leading to their activation and proliferation and differentiation to effector CD8^+^ T cells inducing brain endothelial apoptosis. These observations suggest that during neuroinflammation activated BBB endothelial cells may take up CNS derived Ags from their abluminal side and subsequently cross-present these Ags on their luminal surface in MHC class I to circulating CD8^+^ T cells. It was in fact previously proposed that recognition of their cognate antigen on MHC class I would facilitate CD8^+^ T cell entry into the CNS across the BBB^10^.

T cells circulating in the blood stream are however exposed to shear forces and thus recognition of their cognate Ag on the BBB would require productive engagement of the TCR with the endothelial peptide-MHC class I complex under physiological flow. *In vitro* and *in vivo* live cell imaging has provided evidence that T cell extravasation across the inflamed BBB is a multi-step process regulated by the sequential interaction of adhesion and signaling molecules between the endothelial cells and the T cells (summarized in^8^). We have previously shown that activated CD8^+^ T cells can cross pMBMECs under physiological flow *in vitro* in an antigen independent manner by LFA-1 mediated shear resistant arrest on endothelial ICAM-1 and ICAM-2 followed by their polarization and crawling and finally their G-protein receptor (GPCR)-dependent diapedesis across the pMBMEC monolayer preferentially via transcellular pores^25^. Investigating the interaction of naïve and effector CD8^+^ T cells with the inflamed pMBMECs under physiological flow we here found that effector OT-I cells arrested on the stimulated pMBMECs with significantly higher efficiency than naïve cells. Endothelial Ag-presentation did not affect the shear resistant arrest of naïve nor the effector OT-I cells suggesting that TCR engagement of peptide-MHC molecules on the endothelial surface does not occur under physiological flow. While following their shear resistant arrest the majority of naïve but not effector OT-I cells readily detached from the pMBMEC monolayers, detachment of naïve OT-I cells was prohibited in presence of their cognate Ag and MHC class I. This observation underscores that during transient arrest on pMBMECs naïve CD8^+^ T cells can engage with their TCR endothelial peptide-MHC class I complexes leading to increased avidity of their binding to the brain endothelium probably via inside-out activation of their integrins allowing for high affinity binding of their endothelial ligands^26^. Although endothelial Ag-presentation allowed for adhesion strengthening of naïve CD8 T cells to the pMBMEC monolayer under flow it abrogated T-cell crawling on the pMBMECs in line with previous observations showing that engagement of the TRC with the MHC-peptide complexes induces a stop signal for the T cells and hinders their crawling^27^. Accordingly, we observed that Ag-presentation by inflamed pMBMECs also induced such a stop signal for effector OT-I cells by abrogating their crawling and subsequent diapedesis across the pMBMEC monolayer. Thus, provided their productive initial engagement on brain endothelial cells under physiological shear, naïve and effector CD8 T cells can therefore in a second step recognize their cognate Ag in an MHC class I dependent manner on brain endothelial cells. Unlike previously proposed Ag-recognition on brain endothelial cells did however not facilitate CD8^+^ T-cell diapedesis across the BBB but in contrast rather triggered their arrest and increased their probing behavior on the luminal surface of the BBB. These arrested and probing CD8^+^T cells induced endothelial apoptosis and breakdown of the pMBMEC monolayer under physiological flow *in vitro.* Our observations therefore suggest that luminal MHC class I restricted Ag presentation by the BBB rather than facilitating CD8^+^ T cell migration across the BBB will arrest CD8^+^ T cells on the luminal side of the BBB eventually triggering focal BBB breakdown as observed in MS.

In MS, demyelination, oligodendrocyte death and axonal loss are the major consequences of chronic neuroinflammation. As our *in vitro* observations showed that pMBMECs were able to cross-present Ag provided from their abluminal side on their luminal surface to CD8^+^ T cells we also aimed to understand if cross-presentation of CNS antigens at the luminal BBB can be observed *in vivo.* To this end we made use of ODC-OVA mice expressing ovalbumin as a sequestered CNS model antigen in the cytoplasm of their oligodendrocytes^16^. Upon infection with LCMV-OVA, peripherally activated OT-I cells enter the CNS parenchyma of ODC-OVA but not WT control mice leading to neuroinflammation, oligodendrocyte damage and clinical disease^28,29^. As infection of endothelial cells with the LCMV strain used in our present study has not been observed^30^ and virus is readily cleared from the mouse^17^, we argued that brain endothelial MHC class I dependent cross-presentation of oligodendrocyte derived OVA should lead to increased arrest and reduced crawling of OT-I cells in CNS microvessels in ODC-OVA mice when compared to control mice *in vivo.* Supporting this notion, we indeed observed increased arrest of both, naïve and effector OT-I cells in spinal cord microvessels of ODC-OVA mice when compared to control mice with effector OT-I cells showing more efficient and sustained interactions. Recapitulating our *in vitro* observations, we also observed a significant reduction of effector OT-I cell crawling and sustained interaction in the spinal cord microvessels of ODC-OVA mice when compared to WT control mice.

In summary, we show that brain microvascular endothelial cells can process Ags from their luminal and abluminal side and (cross)-present these Ags on their luminal surface on MHC class I molecules. Unlike expected^10^ rather than facilitating CD8^+^ T cell entry into the CNS, MHC class I restricted Ag-presentation by the BBB endothelial cells was found to arrest CD8^+^ T cells at the luminal BBB and induce brain endothelial apoptosis. Our observations are in line with a previous study where in a transgenic mouse model the targeted expression of a neo-antigen in brain endothelial cells resulted in BBB breakdown upon transfer of neo-antigen specific effector CD8^+^ T cells^31^. The density of peptide-MHC class I complexes on the luminal BBB may thus be decisive for the degree of damage that can be induced by circulating Ag-specific CD8^+^ T cells. That in turn depends on the expression levels of endothelial MHC class I molecules, which are upregulated during inflammation but also the amount and the nature of CNS tissue damage, which will influence Ag processing and presentation by brain endothelial cells. Importantly, we found that the initial adhesive interactions of CD8^+^ T cells with brain microvascular endothelial cells under shear are independent of Ag-recognition. Thus, only those CD8^+^ T cells that are able to initiate productive adhesive interactions with the BBB will be able to subsequently recognize their cognate Ag on the BBB. In the absence of neuroinflammation adhesion molecule expression on BBB endothelium is very low and T cell interaction is restricted to low numbers of activated T cells^8^ suggesting that Ag recognition by CD8^+^ T cells on the BBB rarely occurs under these conditions. Increased adhesion molecule expression during neuroinflammatory diseases allows for increased adhesive interactions mediated by adhesion molecules like α4β1-integrin (VLA-4) which also mediates CD8^+^ T cell entry into the CNS^32^. It is tempting to speculate that the success of therapeutic targeting of this molecule for the treatment of MS not only relies on inhibition of T cell migration across the BBB but in parallel on prohibiting Ag-specific CD8^+^ T cell interactions with the brain endothelium and thus CD8^+^ T cell mediated BBB breakdown. Future studies on how the nature of the respective Ag, the density of Ag presentation on MHC class I molecules on brain endothelial cells affects the multi-step extravasation of CD8^+^ T cells across the BBB and BBB integrity will be relevant to improve our understanding of the pathology underlying MS and other neuroinflammatory disorders.

## Material & Methods

### Mice

Wildtype C57BL/6J mice were obtained from Janvier (Genest Saint Isle, France). β_2_-microglobulin^-/-^ (B6.129P2-B2m^tm1Jae^) mice lacking the functional expression of MHC class I^33^ were provided by Swiss Immunological Mouse Repository (SWImMR; Zürich). OT-I mice (C57BL/6J-Tg(Tcra/Tcrb)1000Mjb) contain a transgenic TCR, recognizing OVA residues 257-264 (SIINFEKL) in the context of H2K^b 22^. OT-I Rag-1^-/-^ C57BL/6J mice were generated by crossing OT-I mice with Rag-1^-/-^ mice (Rag1^tm1Mom^). OT-I tdTomato mice expressing tdTomato reporter ubiquitously were generated by crossing OT-I mice with Ai14 tdTomato reporter mice (Gt(ROSA)26Sor^tm14(CAG-tdTomato)Hze^), in which the Stop-cassette had previously been deleted by breeding with ZP3-Cre transgenic (Tg(Zp3-cre)93Knw) mice. Mice expressing a VE-Cadherin C-terminal EGFP-fusion protein from the endogenous locus (Cdh5^tm9Dvst^) have been described before (^20,34,35^). NG2-DsRed (Tg(Cspg4-DsRed.T1)1Akik mice^36^ were provided by Ralf Adams (MPI, Münster). OT-I Granzyme B^-/-^ mice were generated by crossing OT-I mice with Granzyme B^-/-^ (Gzmb^tm1Ley^) mice^37^ kindly provided by Charaf Benarafa (IVI, Mittelhäusern, Switzerland). ODC-OVA mice have been described before^16^. All gene targeted mice were backcrossed to the C57BL/6J background for at least 10 generations. All mice were housed in individually ventilated cages under specific pathogen-free conditions at 22 °C with free access to chow and water. Animal procedures were approved by the Veterinary Office of the Canton Bern (permit no. BE31/17 and BE55/20) and are in line with institutional and standard protocols for the care and use of laboratory animals in Switzerland.

### *In vitro* blood-brain barrier model

Primary mouse brain microvascular endothelial cells (pMBMECs) were isolated from 7-10 weeks old β2-microglobulin^-/-^ and their C57BL/6J WT littermates or from VE-Cadherin GFP knock in C57BL/6J mice and cultured exactly as previously described^38,39^. Intact monolayers were stimulated with recombinant mouse TNF-α (5ng/mL, Vitaris AG, Baar, Switzerland) and recombinant mouse IFN-γ (100 U/mL, PreproTech EC Ltd., London, UK) 24 hours prior to the assays.

### CD8^+^ T-cell isolation and differentiation

*Naïve CD8^+^ T-cell isolation:* Peripheral lymph nodes and spleens from Rag-1^-/-^ OT-I, OT-I or tdTomato OT-I C57BL/6J mice were harvested and single cell suspensions were obtained by homogenization and filtration through a sterile 100 μm nylon mesh. A second filtration was applied after erythrocyte lysis (0.83% NH4Cl, Tris-HCl). OT-I cells were isolated with magnetic CD8^+^ T cell selection beads (EasySep, STEMCELL Technologies). The purity of the CD8^+^ T cells was assessed by flow cytometry and was >98.5% in each experiment.

*In vitro activation of naïve CD8^+^ T cells:* OT-I CD8^+^ T cells were isolated and activated from OT-I, tdTomato OT-I, Perforin^-/-^ OT-I or Granzyme B^-/-^ OT-I mice exactly as described before^25,40^. Activated T cells were cultured in IL-2 containing media for 3 days post-activation.

*In vivo activation of naïve CD8^+^ T cells:* 2×10^5^ naïve td-Tomato OT-I cells were intravenously (i.v.) injected into WT C57BL/6J mice 24 hours prior to intraperitoneal (i.p) infection with 10^5^ plaque-forming unit (PFU), OVA expressing lymphocytic choriomeningitis virus (LCMV-OVA)^17^. Spleens of recipient mice were collected 8 days after viral infection and CD8^+^ T cells were purified by magnetic bead selection (EasySep, STEMCELL Technologies). Fluorescence-activated cell sorting is used for the separation of tdTomato OT-I CD8^+^ cells from the CD8^+^ T cells of recipient WT mice.

### Bone marrow derived dendritic cell isolation and culture

Bone marrow derived dendritic cells (BMDCs) were isolated from WT C57BL/6J mice as described before ^41^. In brief, following removal of the femurs and tibiae, the tips of these bones were cut open and the BM cells were isolated by centrifugation (4 min, 1500g). Collected BM cells were cultured untouched in 20 ml cultures in petri dishes containing 18 ml restimulation medium (RPMI-1640 supplemented with 10% FBS (Thermo Fisher Scientific), 10 U/ml penicillin-streptomycin, 2 mM L-glutamine, 1% non-essential amino acids, 1 mM sodium pyruvate, and 0.05 mM β-mercaptoethanol (Grogg Chemie AG) and 2 ml Flt-3L-containing supernatant [produced from SP2/0 transfected cell line secreting mouse recombinant Flt-3L^42^ for 7 days at 37°C until activation with 100 ng/mL LPS (Sigma-Aldrich) 18 hours prior to the assays.

### Immunofluorescence staining of pMBMECs

Confluent pMBMEC monolayers were stained as described before^39^. Primary and secondary antibodies are listed in Supplementary Table 1. The assessment of apoptosis in pMBMECs was performed by immunofluorescence staining with the Image-iT™ LIVE Red Poly Caspases Detection Kit (Thermo Fisher Scientific, Massachusetts, USA) according to the manufacturer’s protocol. As a positive control, apoptosis of pMBMECs was induced by incubation with staurosporine (1μM; Abcam, Cambridge, UK). Images were acquired using a Nikon Eclipse E600 microscope connected to a Nikon Digital Camera DXM1200F with the Nikon NIS-Elements BR3.10 software (Nikon, Egg, Switzerland) or an LSM 800 (Carl Zeiss, Oberkochen, Germany) confocal microscope. Images were processed by using ImageJ software (ImageJ software, National Institute of Health, Bethesda, USA) and mounted in Adobe Illustrator software.

### Quantitative Real-Time PCR (qRT-PCR)

Sample RNA extraction was performed from freshly isolated C57BL/6J WT pMBEMCs following the manufacturers instruction of the High Pure RNA Isolation Kit (Hoffman-La Roche). cDNA was obtained from total isolated RNA of each sample with the SuperScript III First-Strand Synthesis System (Invitrogen, Carlsbad, CA, USA). mRNA expression was analyzed exactly as described before^43^. Beta actin mRNA levels were used as endogenous control. The sequences of the primers used for each gene are presented in Supplementary Table 2.

### Flow cytometry

Cell surface molecules of T cells and DCs were stained with appropriate combinations of fluorophore-conjugated mAbs at saturating concentrations on ice in the dark for 30 min. Intracellular staining of T cells was performed exactly as described before^41^. The antibodies used and the working concentrations are listed in Supplementary Table 1. The data was acquired using an Attune NxT flow cytometer (ThermoFisher Scientific, Massachusetts, USA) and analyzed using FlowJo 10 software.

### Co-culture assays of naïve CD8^+^ T cells with pMBMECs/BMDCs

The activation of naïve OT-I cells was assessed by flow cytometry. TNF-α/IFN-γ-stimulated WT or B2M^-/-^ pMBMECs were pulsed on the luminal side in 96-well format with either SIINFEKL (POV-3659-Peptides International, Kentucky, USA) or the (VSV) nucleoprotein peptide (RGYVYQGL, BACHEM, Bubendorf, Switzerland) at 0.1 ng/mL for 30 minutes at 37°C or left unpulsed. The VSV)-peptide binds to H-2K^b^ but is not recognized by the TCR of the OT-I cells^44^ and was therefore used as a control.

Abluminal assays were performed by employing Transwell system (0.4 um pore size, 0.33 cm^2^ filter inserts: Transwell^®^, Costar, Corning). SIINFEKL, VSV-peptide and endotoxin free-, full length ovalbumin (OVA; EndoFit, InvivoGen, San Diego, CA, USA) protein were loaded in the pMBMECs with overnight incubation at 37°C. Unprocessed peptides were removed by 3x wash with wash buffer containing 10mM HEPES and 0.1% BSA. Consecutively, 5×10^5^ naïve OT-I cells per well were co-incubated with pMBMECs. Professional Ag presenter, LPS-stimulated BMDCs were co-incubated with the naive OT-I cells 1:3 ratio (BMDCs/CD8^+^ T cells) as a positive control for luminal assays. The naïve status of the cells was evaluated before co-incubation by flow cytometry staining with CD25, CD69, CD44 and homing receptor CD62L (Data not shown; Antibodies: Supplementary Table 1). After 24 hours, the T cells were collected from pMBMEC- or BMDC co-cultures and were stained for the same markers for the detection of activation. By employing the exact co-culture system, the induction of proliferation of naïve CD8^+^ T cells was assessed by BrdU Cell proliferation ELISA kit (Abcam, Cambridge, UK) according to the protocol of the manufacturer after 72 hours of co-culture. Following the removal of the T-cells, pMBMECs were washed and immunofluorescent stained with ZO-1 and DAPI. As an internal control, pMBMECs that were pulsed with peptides but not co-cultured with CD8^+^ T cells were stained with the same markers to control the T-cell unrelated effects.

### *In vitro* live-cell imaging

*In vitro* live cell imaging of T-cell interactions with pMBMECs was performed as described before^39^,^20^,^45^. Briefly, prior to the assay WT and B2M^-/-^ pMBMECs are pulsed with either SIINFEKL or VSV peptide (0.5 ng/mL) for 30 minutes at 37°C or left unpulsed. Following this step, they were gently washed 3x with wash buffer C (1x HBSS with Ca^2+^ Mg^2+^ (Gibco), 10mM HEPES (Gibco), 0.1% BSA) to remove the unprocessed peptide. OT-I or tdTomato OT-I CD8^+^ T cells (1×10^6^ cells/ml) were superfused over WT or B2M^-/-^ pMBMECs. Accumulation of CD8^+^ T cells on pMBMECs in the flow chamber was allowed at a low shear (0.1 dyn/cm^2^) for 5 minutes, followed by physiological shear (1.5 dyn/cm^2^) for an additional 25 minutes to assess the post-arrest behavior or 55 minutes to observe the late dynamics. Images were acquired at 10x or 20x magnification with an inverted microscope (AxioObserver, Carl Zeiss, Oberkochen, Germany) with phase-contrast and fluorescence illumination, every 10 seconds. Image analysis was performed using ImageJ software (ImageJ software, National Institute of Health, Bethesda, USA). Thirty seconds after the onset of the enhanced shear, the number of arrested T cells were counted manually by using ImageJ. The post-arrest behavior of T cells was defined and visualized as fractions of categorized T cells set to 100%, as follows: T cells that actively sent protrusions underneath the endothelium in a stationary position (probing); polarized T cells that continuously crawled on the endothelium (crawling); T cells that transmigrated across the endothelium following probing or crawling (probing + diapedesis, crawling + diapedesis); T cells that did not complete a diapedesis event with prior probing or crawling (probing + partial diapedesis and crawling + partial diapedesis); T cells that detached during the observation time (detaching). T cells that crawled out of the imaged field of view or went through cell division were counted as arrested but not categorized for the behavior. T-cell crawling tracks were evaluated after manual tracking of individual T cells, using the manual tracking plug-in of ImageJ.

### CD8^+^ T-cell-mediated CNS autoimmune disease: ODC-OVA Model

CD8^+^ T cell mediated autoimmune disease was induced in 9–12-week-old ODC-OVA mice by intravenous injection of freshly isolated naïve tdTomato OT-I cells (2×10^5^ cells in 100 μl/mouse) and wild-type C57BL/6J mice were used as controls. 24 hours later, mice were peripherally challenged with an intraperitoneal injection of 10^5^ plaque-forming units (PFU) of LCMV-OVA^17^. Animals were monitored twice per day for clinical symptoms and scored as follows: 0, healthy; 0.5, limp tail; 1, hind leg paraparesis; 2, hind leg paraplegia; 3, hind leg paraplegia with incontinence as previously described^17^.

### Intravital microscopy of the cervical spinal cord

Epifluorescence and 2PM intravital microscopy of cervical spinal cord of WT, VE-Cadherin-GFP knock in and ODC-OVA transgenic C57BL/6J mice was performed exactly as described before^40^,^46^. Both, naïve and activated OT-I cells were stained with 1 uM CellTracker Green CMFDA dye (Thermo Fisher Scientific) and systemically injected via a carotid artery catheter as described^46^. For 2PM imaging, the blood vessels were *in vivo* stained with Alexa Fluor 633-conjugated rat-anti mouse endoglin antibody (20 μg/mouse) as described before^46^. Distortion correction of the images during 2PM-imaging was performed by using Vivo Follow 2.0^47^. Sequences of image stacks were transformed into volume-rendered 4D images by Imaris 9.2 software.

### Statistical analysis

Statistical analysis was performed using GraphPad Prism 7.0 software (La Jolla, CA, USA). Data were compared using various tests as indicated in each figure legend. Data are presented as mean ±SD or ±SEM and asterisks indicate significant differences (\**p* < 0.05, \*\**p* < 0.01 and \*\*\**p* < 0.001, \*\*\*\**p* < 0.0001).

## Supporting information

Supplementary Video 1. Priming of naive OT-I cells by pMBMECs upon endothelial Ag-presentation.

Supplementary Video 2. Naive OT-I T-cell interactions with pMBMECs under physiological flow

Supplementary Video 3. Effector OT-I T-cell interactions with pMBMECs under physiological flow

Supplementary Video 4. OT-I T cell induced apoptosis of Ag-presenting WT pMBMECs under physiological flow

Supplementary Video 5. OT-I vs OT-I GrB-/- T cell interactions with VE-Cadherin-GFP pMBMECs under physiological flow

Supplementary Video 6. OT-I CD8+ T cell probing on the BBB under neuroinflammation

Supplementary Video 7. OT-I CD8+ T cell probing and crawling on the BBB

## Acknowledgements & Funding

We thank to Dr. Gaby Enzmann her support with the biosafety level 2 work. Additional thanks go to Charaf Benarafa (IVI, Mittelhäusern, Switzerland) for providing the Granzyme B^-/-^ C57BL/6 mice and to Thomas Hünig (Institute of Virology and Immunobiology, Würzburg, Germany) for providing the ODC-OVA C56BL/J mice. This project has been funded by the Swiss National Science Foundation (grant numbers 31003A_149420 and 310030_189080) and the Fondation Pour L’Aide a la Recherche sur la Sclérose en Plaques (ARSEP). DM is supported by the Swiss National Science Foundation (grant numbers 310030B_201271 and 310030_185321) and ERC. AAJ is supported by NIH grants R01 NS103212 and RF1 NS122174.

## Contributions

SA designed, performed and analyzed all the experiments and wrote the manuscript. BE designed the overall study and the experiments, participated in data analysis and wrote the manuscript. JP performed and analyzed experiments and contributed to manuscript writing. VMS, AK and TG performed and analyzed experiments. EK performed experiments and quality control. UD managed the mouse colonies. NP, DM, AJJ, MS provided reagents, cells or mice and methods and advised in experimental design.

