## Supplementary Documents for "Brain endothelial antigen presentation detains CD8^+^ T cells at the blood-brain barrier leading to its breakdown"

[1]Theodor Kocher Institute, University of Bern, Bern, Switzerland, [2]Institute of Pathology, Experimental Pathology, University of Bern, Bern, Switzerland, [3]Department of Pathology and Immunology, Division of Clinical Pathology, University and University Hospitals of Geneva, Geneva, Switzerland, [4]Mayo Clinic Graduate School of Biomedical Sciences, College of Medicine, Mayo Clinic, Rochester, MN, USA.

**Keywords:** $CD8^+$ T cells, antigen-presentation, blood-brain barrier, ODC-OVA mouse, multiple sclerosis

**Running Title:** Brain endothelial antigen presentation prohibits $CD8^+$ T cell diapedesis

**Corresponding author:**
Prof. Dr. Britta Engelhardt
Theodor Kocher Institute
University of Bern
Freiestrasse 1
CH-3012 Bern
Switzerland
ORCID:orcid.org/0000-0003-3059-9846

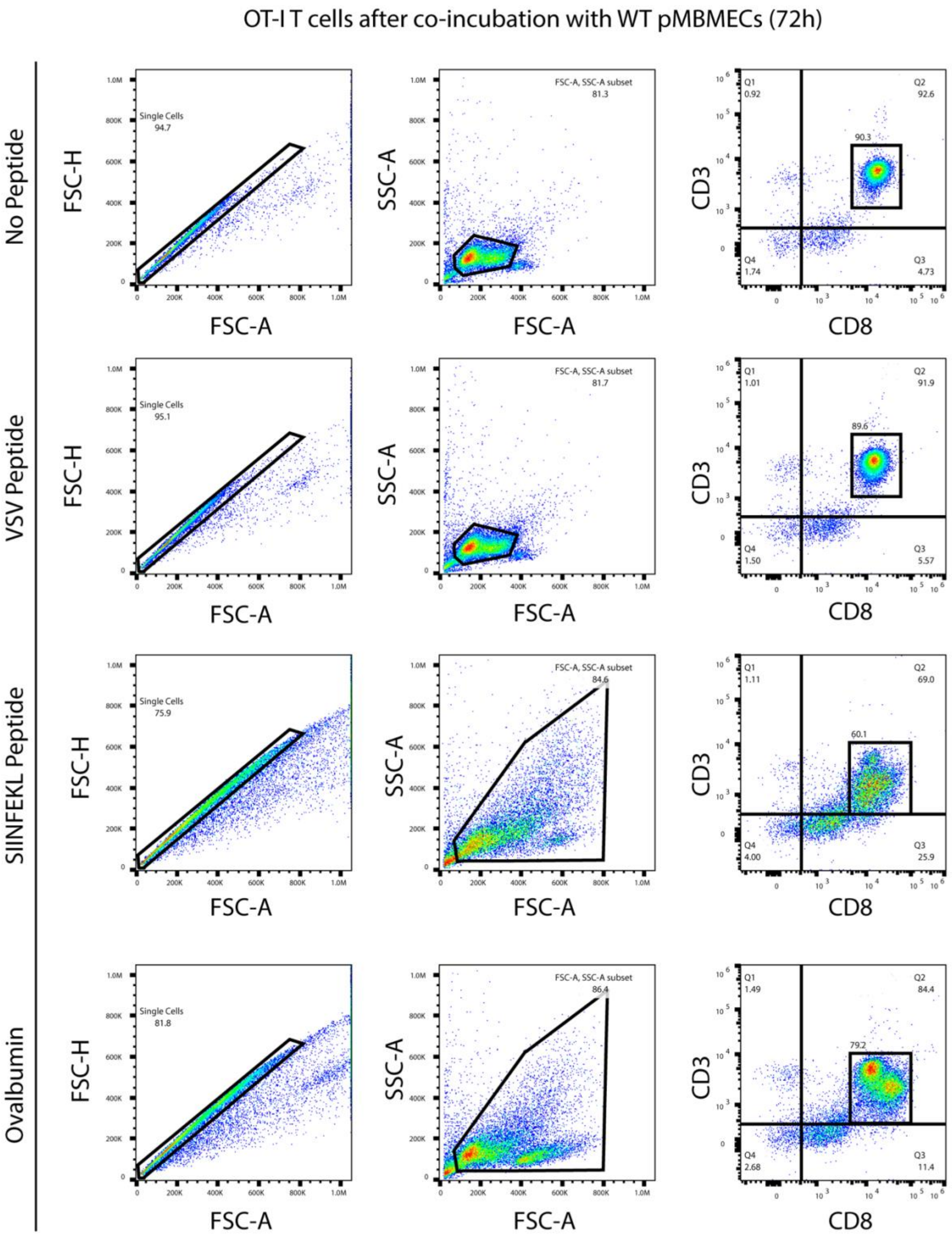


**Supplementary Figure 1. Gating strategy for OT-I cells upon co-culture with WT pMBMECs**

Flow cytometry analysis of naïve OT-I cells after 24 hours of co-culture with TNF-$\alpha$/IFN-$\gamma$ stimulated WT pMBMECs in the absence of antigen, or in the presence of VSV- or SIINFEKL peptide or full-length ovalbumin (OVA) protein. Single cells are selected first by using forward scatter height versus forward scatter area (FSC-H::FSC-A). $CD3^+CD8^+$ cells were selected from the population of interest based on the side scatter area versus forward scatter area (SSC-A::FSC-A). OT-I $CD8^+$ T cells that are co-cultured with unpulsed or VSV-pulsed pMBMECs did not show an increase in their size and granulation, whereas SIINFEKL pulsed- or OVA-loaded pMBMECs induced a major increase in the size and granulation of the naïve OT-I $CD8^+$ T cells, indicating their activation. Moreover, activated OT-I $CD8^+$ T cells mostly lost their $CD3^+$ cell surface expression.

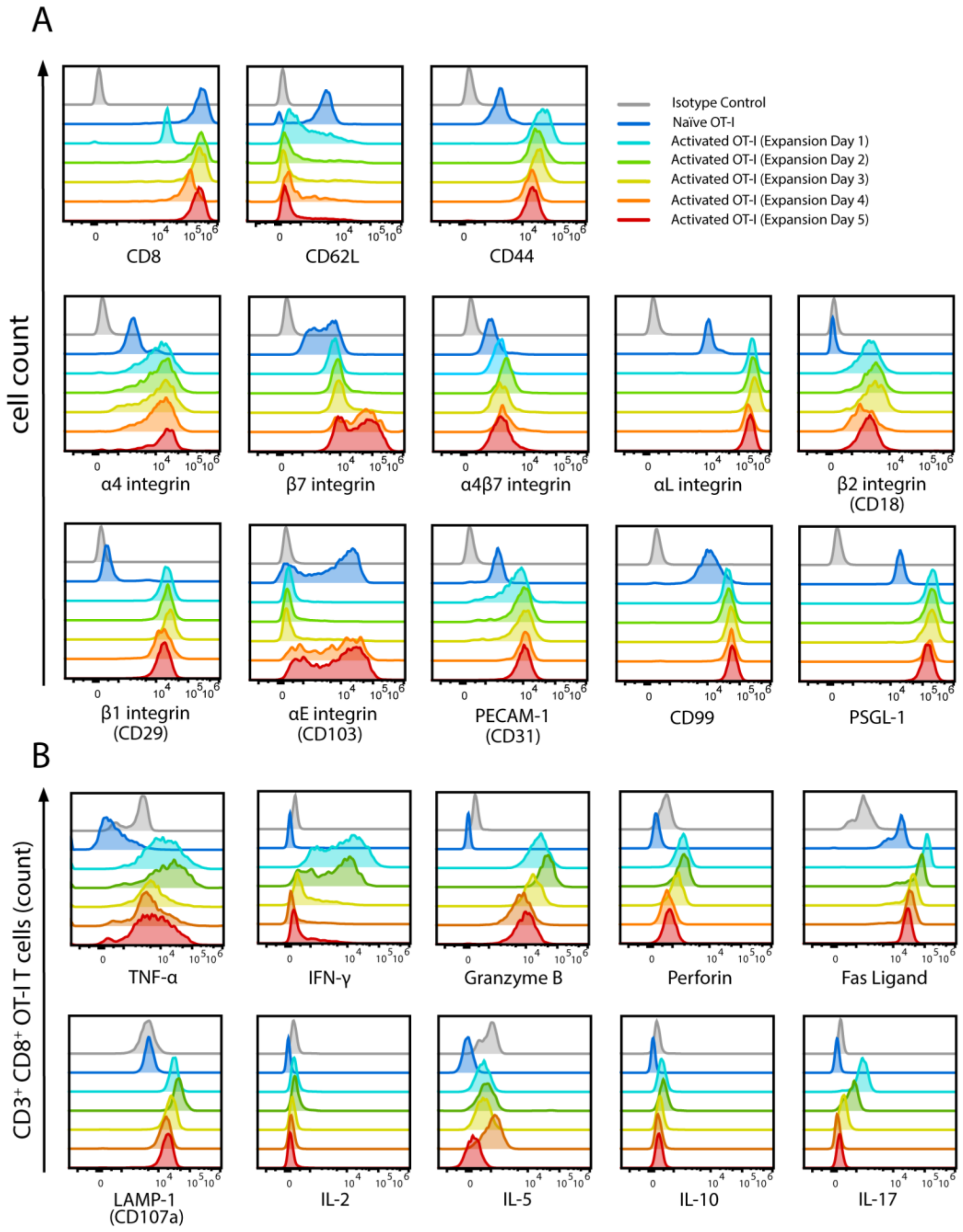


**Supplementary Figure 2. Phenotype of *in vitro* activated OT-I cells**

Phenotpy of *in vitro* activated and expanded OT-I cells by flow cytometry at different stages of their expansion. Naïve OT-I cells were used to compare the changes following the *in vitro* activation. **(A)** Single color flow cytometry staining of OT-I cells for activation and cell surface expression of T cell trafficking related markers; CD8, CD44, CD62L, α4-, ß7-, α4ß7-, αL-, ß2-, αE-, ß1-integrins, PECAM-1, CD99, PSGL-1. At the expansion day 1, OT-I cells are not fully activated based on the loss of cell surface staining for CD62L, while at the expansion day

4 and 5, OT-I T cells show the upregulation of αE-integrin, indicating a tissue memory cell phenotype. **(B)** Flow cytometry staining of $CD3^+CD8^+$ OT-I cells for cytotoxicity related molecules granzyme B, perforin, Fas ligand, LAMP-1 and cytokines TNF-α, IFN-γ, IL-2, IL-5, IL-10, IL-17.

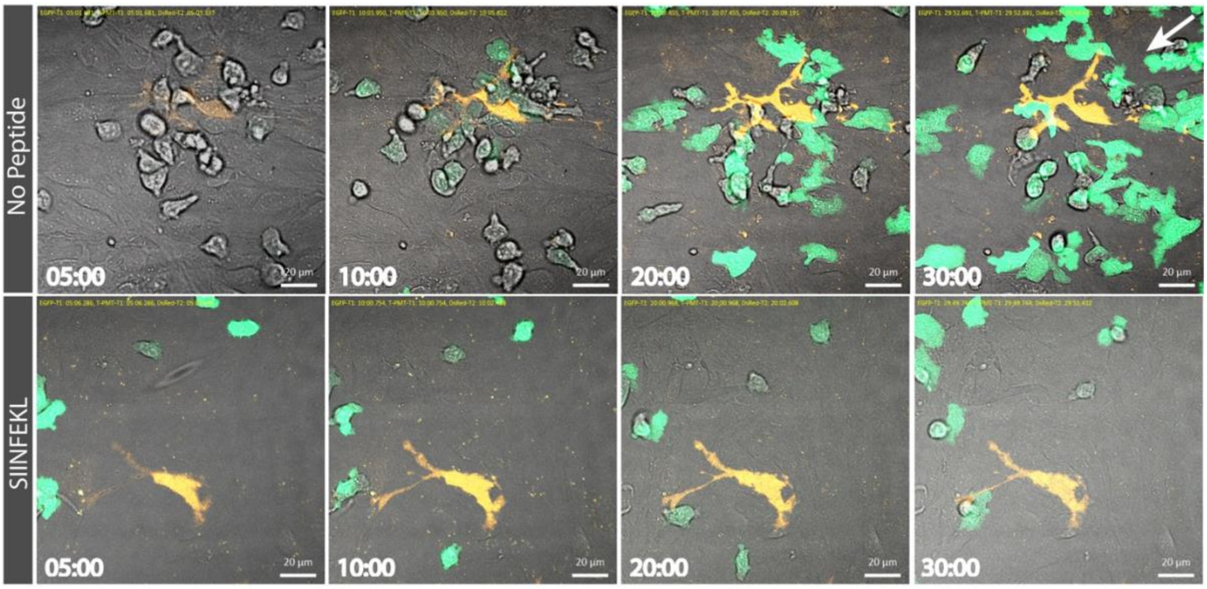


**Supplementary Figure 3. Pericytes do not contribute to Ag-presentation to OT-I cells in pMBMEC monolayers under flow**

Interaction of CMFDA-labeled activated OT-I T cells with TNF-α/IFN-γ stimulated pMBMECs from NG2- DsRed C57BL/6J mice pulsed or not with SIINFEKL peptide at different timepoints under physiological flow is shown. $1x10^6$ OT-I cells were superfused over the pMBMECs. Images are recorded for 30 minutes with LSM 800 confocal microscope (Carl Zeiss, Oberkochen, Germany). White arrow indicates the direction of the flow. OT-I T cells (green) do not specifically interact with the pericyte (red) in the SIINFEKL-pulsed pMBMEC monolayer. Scale bar = 20 µm.

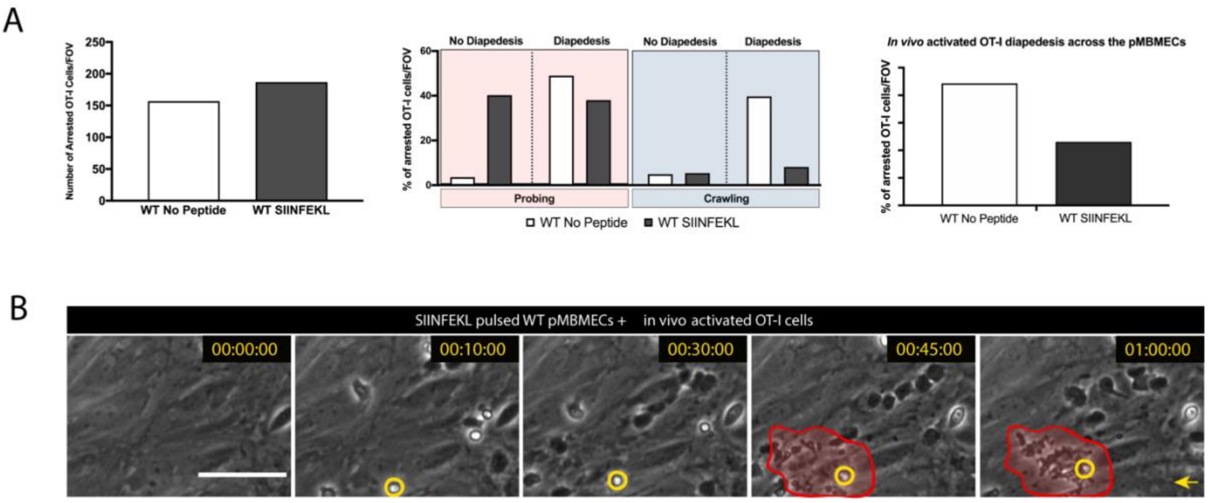


**Supplementary Figure 4. Interaction of *in vivo* activated OT-I CD8$^+$ T cells with pMBMECs compares to that of *in vitro* activated OT-I CD8$^+$ T cells.**

*In vivo* activation of tdTomato$^+$ naïve OT-I cells was performed by their injection into a WT C57BL/6J mice prior to LCMV-OVA infection. Spleens of the recipient mice were collected at day 8 post-infection and single cells were purified by magnetic bead selection and fluorescence-activated cell sorting. **(A)** Bar graphs show the dynamic interaction of *in vivo* activated OT-I cells with TNF-$\alpha$/IFN-$\gamma$ stimulated pMBMECs under physiological flow *in vitro*. Number of arrested *in vivo* activated OT-I cells on unpulsed or SIINFEKL-pulsed pMBMECs is shown on the left, while the bar graphs in the middle and on the right depict their post-arrest behavior. Data from one experiment are shown. **(B)** Representative image sequence of *in vivo* activated- effector OT-I cell induced killing of SIINFEKL-pulsed pMBEMCs under physiological flow during 60 minutes of imaging. One OT-I CD8$^+$ T cell is highlighted by a yellow circle, while the area of pMBMEC killing is marked with the red area. The yellow arrow indicates the direction of the flow. Scale bar = 20 $\mu$m.

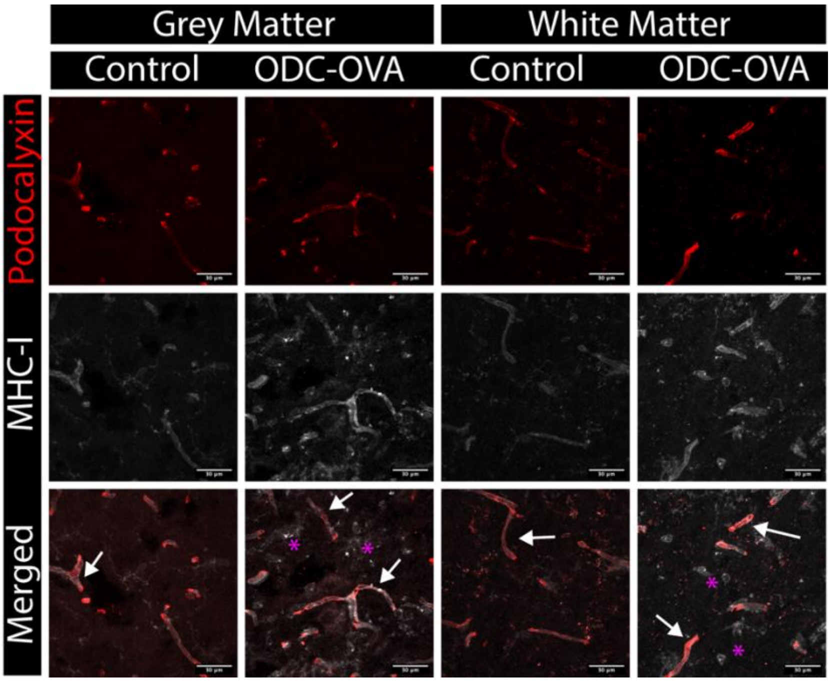


**Supplementary Figure 5. MHC-I colocalization with CNS vasculature in the spinal cord of ODC-OVA and WT C57BL/6 mice.**

Immunofluorescence staining of WT C57BL/6J and ODC-OVA mice spinal cords on day 7 after LCMV-OVA infection for MHC class I (grey), and endothelial cells (podocalyxin, red). Stainings in grey matter are displayed in the left panel and in the white matter on the right panel. White arrows show CNS vessels with a signal from both MHC class I and podocalyxin. Purple asterisks point to other CNS resident cells staining positive for MHC class I molecules on their surface. Scale bar = 30 μm

| Flow Cytometry Antibodies | | | |
|---|---|---|---|
| **Primary Antibodies** | **Company** | **CAT Nr.** | **Working Conc.** |
| Rat monoclonal anti - Mouse CD25 - FITC | BioLegend | 102006 | 5 µg/mL |
| Armenian Hamster monoclonal anti- Mouse CD3 - APC | BioLegend | 100312 | 5 µg/mL |
| Armenian Hamster monoclonal anti - Mouse CD3 - BV605 | BioLegend | 100351 | 8 µg/mL |
| Rat monoclonal anti - Mouse CD44 - eFluor450 | Thermo Fisher Scientific | 48-0441-80 | 5 µg/mL |
| Rat monoclonal anti - Mouse CD62L - PE | BioLegend | 104408 | 0.2 µg/mL |
| Armenian Hamster monoclonal anti - Mouse CD69 - BV711 | BioLegend | 104537 | 4 µg/mL |
| Rat monoclonal anti - Mouse CD8 - PE-Cy7 | BioLegend | 100722 | 6 µg/mL |
| Rat monoclonal - Mouse CD8 - PerCP | BioLegend | 100732 | 2 µg/mL |
| Mouse monoclonal anti - Mouse FasL - PE | BioLegend | 106805 | 10 µg/mL |
| Mouse monoclonal anti - Mouse Granzyme B - FITC | BioLegend | 515403 | 10 µg/mL |
| Rat monoclonal anti - Mouse IFN-gamma - BV711 | BioLegend | 505836 | 10 µg/mL |
| Rat monoclonal anti - Mouse LAMP-1 (CD107a) - PerCP-eFluor710 | Thermo Fisher Scientific | 46-1071-80 | 10 µg/mL |
| Rat monoclonal anti - Mouse Perforin - APC | Thermo Fisher Scientific | 17-9392-80 | 10 µg/mL |
| Rat monoclonal - Mouse TNF-alpha - BV510 | BD Biosciences | 563386 | Variant |
| Monoclonal Armenian Hamster IgG - BV605 | BioLegend | 400944 | Variable |
| Monoclonal Rat IgG2a - PE-Cy7 | Thermo Fisher Scientific | 25-4321-81 | Variable |
| Monoclonal Rat IgG1 - BV510 | BD Biosciences | 563039 | Variable |
| Monoclonal Rat IgG1 - BV711 | BioLegend | 400441 | Variable |
| Monoclonal Mouse IgG1 - FITC | BD Biosciences | 554679 | Variable |
| Monoclonal Armenian Hamster IgG - APC | BioLegend | 400911 | Variable |
| Monoclonal Rat IgG2b - eFluor450 | Thermo Fisher Scientific | 48-4031-80 | Variable |
| Monoclonal Rat IgG2a - PE | BioLegend | 400508 | Variable |
| Monoclonal Rat IgG2a - PerCP | BioLegend | 400530 | Variable |
| Monoclonal Mouse IgG2b - PE | BioLegend | 400314 | Variable |
| Monoclonal Armenian Hamster IgG - BV711 | BioLegend | 400963 | Variable |
| Monoclonal Rat IgG2a - PerCP - eFluor710 | Thermo Fisher Scientific | 46-4321-80 | Variable |
| Monoclonal Rat IgG2a - APC | BioLegend | 400512 | Variable |

| Immunofluorescence Staining Antibodies | | | |
|---|---|---|---|
| **Primary Antibodies** | **Company** | **CAT Nr.** | **Working Conc.** |
| Rat monoclonal anti - Mouse CD8 | BioLegend | 100702 | 10 µg/mL |
| Armenian Hamster monoclonal anti - Mouse CD80 | BioLegend | 104702 | 10 µg/mL |
| Rat monoclonal anti - Mouse CD86 | BioLegend | 105002 | 10 µg/mL |
| Rat monoclonal anti - Mouse MHC class I | BMA Biomedicals | T-2105 | 20 µg/mL |
| Rat monoclonal anti - Mouse PD-L1 | BioLegend | 124302 | 2.0 µg/mL |
| Goat polyclonal anti - Mouse Podocalyxin | R&D Systems | AF1556 | 2.0 µg/mL |
| Rabbit polyclonal anti - Mouse ZO-1 | Thermo Fisher Scientific | 61-7300 | 1.25 µg/mL |
| Rat monoclonal anti - Rat IgG2a | BioLegend | 400501 | Variable |
| Monoclonal Armenian Hamster IgG | BioLegend | 400901 | Variable |
| Monoclonal Rat IgG2b | BioLegend | 400601 | Variable |
| Polyclonal Rabbit IgG | R&D Systems | AB-105-C | Variable |
| **Secondary Antibodies** | **Company** | **CAT Nr.** | **Working Conc.** |
| Donkey polyclonal anti - Rabbit IgG - AF488 | Thermo Fisher Scientific | A-21206 | 10 µg/mL |
| Goat polyclonal - anti - Rat IgG - Cy3 | Jackson Immuno Research | 112-165-143 | 2.5 µg/mL |
| Goat polyclonal Biotin - anti - Hamster | BioLegend | 405501 | 10 µg/mL |
| Goat polyclonal - anti - Rabbit IgG - AF647 | Thermo Fisher Scientific | A-21244 | 3 µg/mL |
| Goat polyclonal - anti - Goat IgG - Cy3 | Jackson Immuno Research | 705-165-147 | 5 µg/mL |
| Donkey polyclonal - anti-Rat IgG - AF488 | Thermo Fisher Scientific | A-21208 | 5 µg/mL |
| Streptavidin Cy3 | BioLegend | 405215 | 10 µg/mL |

**Supplementary Table 1. Sequence of primers used for the qPCR analysis.**

| Official Gene Name | Entrez Gene ID | Accession Number | Primer Sequences | | Supplier |
|---|---|---|---|---|---|
| CD80 (mouse) | 12519 | NM_009855 | Quantitech primer assay for mouse CD80, QT00129787 | | Qiagen |
| CD86 (mouse) | 12524 | NM_019388 | Quantitech primer assay for mouse CD86, QT01055250 | | Qiagen |
| β-actin (mouse) (endogenous control) | 11461 | NM_007393 | Sense | 5'- CGTGGGCCGCCCTAGGCACCA -3’ | Eurogentec |
| | | | Anti sense | 5'- TTGGCTTAGGGTTCAGGGGGG -3’ | |

**Supplementary Table 2. List of antibodies used in this study.**

**Supplementary Video 1. Priming of naïve OT-I cells by pMBMECs upon endothelial Ag-presentation.**

Time lapse imaging of TNF-$\alpha$/IFN-$\gamma$ stimulated, unpulsed or SIINFEKL-pulsed pMBMECs from VE-cadherin-GFP C57BL/6J mice with tdTomato expressing naïve OT-I $CD8^+$ T cells under static conditions for 72 hours. Images are acquired by using a high content imaging system INCell Analyzer 2000 every 30 minutes. Cytokine stimulated-, peptide pulsed pMBMECs without T cells are used as an internal control. tdTomato expressing OT-I cells that are co-cultured with SIINFEKL-pulsed pMBMECs show an increase in size, indicating their activation and is followed by their proliferation. Activated OT-I cells fully disrupt the pMBMECs monolayer that is indicated by the loss of GFP signal at the junctions after their interactions. On the other hand, naïve OT-I cells that are co-cultured with unpulsed pMBMECs lose their cytoplasmic expression of tdTomato and die over 72 hours, while the pMBMECs remain intact. Cytokine stimulated-, SIINFEKL pulsed pMBMECs remain intact in the absence of $CD8^+$ T cells.

**Supplementary Video 2. Naïve OT-I T-cell interactions with pMBMECs under physiological flow**

Time lapse imaging of TNF-$\alpha$/IFN-$\gamma$ stimulated, VSV- or SIINFEKL-pulsed pMBMECs from WT or $B2M^{-/-}$ C57BL/6J mice with naïve OT-I $CD8^+$ T cells under physiological flow for 30 minutes. Images are acquired with 10 seconds intervals. Different post-arrets behaviors are labeled with different circles (Probing: Green; Crawling: Orange; Detachment: Pink). Yellow arrow indicates the direction of the flow. Scale bar = 50 $\mu$m.

**Supplementary Video 3. Effector OT-I T-cell interactions with pMBMECs under physiological flow**

Time lapse imaging of TNF-$\alpha$/IFN-$\gamma$ stimulated, VSV- or SIINFEKL-pulsed pMBMECs from WT or $B2M^{-/-}$ C57BL/6J mice with *in vitro* activated OT-I $CD8^+$ T cells under physiological flow for 30 minutes. Images are acquired with 10 second intervals. Different post-arrets behaviors are labeled with different circles (Probing: Green; Crawling: Orange; Diapedesis: Blue). Yellow arrow indicates the direction of the flow. Scale bar = 50 $\mu$m.

**Supplementary Video 4. OT-I T cell induced apoptosis of Ag-presenting WT pMBMECs under physiological flow**

Time lapse imaging of TNF-$\alpha$/IFN-$\gamma$ stimulated, SIINFEKL-pulsed pMBMECs from WT C57BL/6J mice with *in vitro* activated OT-I $CD8^+$ T cells under physiological flow for 60 minutes. Images are acquired with 10 second intervals. Yellow arrow indicates the direction of the flow. Scale bar = 50 μm.

**Supplementary Video 5. OT-I vs OT-I $GrB^{-/-}$ T cell interactions with VE-Cadherin-GFP pMBMECs under physiological flow**

Time lapse imaging of TNF-$\alpha$/IFN-$\gamma$ stimulated, unpulsed or SIINFEKL-pulsed pMBMECs from VE-Cadherin GFP C57BL/6J mice with in vitro activated effector OT-I or OT-I $GrB^{-/-}$ $CD8^+$ T cells under physiological flow for 60 minutes. Yellow circle highlights the disruption of the junctions and the pMBMEC-monolayer. Scale bar = 20 μm.

**Supplementary Video 6. OT-I $CD8^+$ T cell probing on the BBB under neuroinflammation**

Time lapse imaging of two-photon intravital microscopy of Cell Tracker Green CMFDA labeled effector OT-I $CD8^+$ T cells (green) adhered in inflamed cervical spinal cord microvessels (white) of ODC-OVA C57BL/6J mice over 30 minutes on day 7 after viral infection. The video is depicted as maximum intensity projection of 100 μm thick Z-stack. Green circle indicates a cell probing. Scale bar = 50 μm

**Supplementary Video 7. OT-I $CD8^+$ T cell probing and crawling on the BBB**

Time lapse imaging of two-photon intravital microscopy of Cell Tracker Green CMFDA labeled effector OT-I $CD8^+$ T cells (green) adhered in inflamed cervical spinal cord microvessels (white) of ODC-OVA C57BL/6J mice over 40 minutes on day 7 after viral infection. The video is depicted as maximum intensity projection of 100 μm thick Z-stack. Different post-arrets behaviors are labeled with different circles (Probing: Green; Crawling: Orange). Scale bar = 50 μm.
